# Off-target prediction of SGLT2 inhibitors: an integrative bioinformatics approach to uncover structural mechanisms

**DOI:** 10.1101/2025.01.20.633821

**Authors:** Hsin-Ju Yang, Yun-Ti Chen, Zi-Chun Hsu, Jinn-Moon Yang

## Abstract

Gliflozin is known to inhibit sodium glucose transporter 2 (SGLT2) inhibitors (SGLT2i). They have been approved for the treatments of diabetes mellitus, cardiovascular diseases, and chronic kidney disease in recent years. However, the mechanisms for the multifunction remain unclear. We here propose a hypothesis of other protein targets for gliflozins. Additionally, the multistage of the protein, the specificity of the drugs, along with the structural data shortage have posed a hindrance to disclose gliflozin and its binding environment (BE).

Considering the difficulties on the issue, this thesis provides an approach to uncover protein off-targets and to find the underlying pathways. We predicted pyranose was a critical substructure from > 1,500 SGLT2i. Hierarchical clustering on atom-based interaction, combined with propensity, revealed key interaction among 1,572 pyranose BEs. The other criterium presented was the concept of compound spanning space via protein pocket size. Binding sites (BSs) from proteins like SGLT2 were able to provide similar pockets, approximating the occupied space of the ligands. Finally, gliflozin binding pockets prefer (1) aromatic residues for van der Waals force, (2) aspartate or asparagine as hydrogen bond providers, and (3) protein pocket size ≧442 Å ^2^. These criteria were integrated into a scoring function S_T_ and predicted several possible proteins.

This research presents a pipeline for target identification of emerging drug agents and proteins with crystallographic difficulties. The study provides an innovative computational methodology on extraction of binding features on transmembrane proteins and further pharmaceutical development.

## Introduction

Diabetes mellitus (DM), cardiovascular disease (CVD), and chronic kidney disease (CKD) are three closely related diseases on the top 10 leading causes of death around the world^1^. The mechanisms causing the intimate association of DM with CKD and CVD have been widely reported^2-4^. In the United States, 69% of adult patients with DM had high blood pressure, while 39% had CKD. Among the 8.27 million hospital discharges for diabetes in 2018, 1.87 million were for CVD^5^. The main claim of gliflozins’ mechanism is that higher glucose levels in blood may lead to raised cardiovascular stress^4^ and renal pressure to filter and produce urine^6^. Hyperglycemia would cause damage to the glomerular system and activate the renin and angiotensin aldosterone system^7^ to increase the work load of heart and kidney^8^. The increasing stress on the systems leads to disease eventually.

Current studies manifest that a type of drug, gliflozins, can treat all three^9-11^. Gliflozins, also known as human sodium glucose transporter 2 (hSGLT2) inhibitors (hSGLT2i), are composed of β-glucopyranose moiety and a tail with two aromatic rings of a distance of two single-bonds^12^. They were derived from a natural compound, phlorizin^13^. Chemical structures were further improved for metabolic stability and inhibitory effects^14^ on hSGLT2, which was the invention of hSGLT2i. Unlike the previous medication for DM rebalancing insulin-missing endocrine signaling pathways^3^, gliflozins ease hyperglycemia (high-level blood sugar) by blocking hSGLT2^15^.

Gliflozins was originally approved as a DM medication^11^. Afterwards, clinicians have noticed the alleviation of complications of CKD or CVD^16-18^. In the past two years, they have also been approved for treatment on limited CVD^9^ and CKD^10^. Ever since the clinical trials reveal the multifunctionality of gliflozins^17-20^, many publications have proposed the potential mechanisms^21-23^. The basis is that by inhibiting hSGLT2, hyperglycemia can be alleviated. Inhibition of hSGLT2 in the kidney and reduction of glucose concentration in patients with DM ease work load on the renal system^21^. Furthermore, controlled glucose levels reduce heart stress. However, studies have revealed that some glucose-lowering effect in other drugs does not improve cardiovascular functions^16,24-27^. For comparisons with patients taking other DM treatments, researchers have seen an increased expression in the glucose transporter type 4 (GLUT4) in epicardial adipose^28^, lowered cytokine secretion and ease of inflammatory effects^29^, and cardiovascular protecting effects^9,10,30^. These all indicate there are still other unclear mechanisms of gliflozins. We thus propose the hypothesis there are other protein targets for gliflozins that lead to multifunctioning and uncovered molecular pathways to treat DM, CVD, and CKD at the same time.

Generally in computational structural biology, drug off-target prediction is based on the concept of Homopharma^31^ that similar ligands bind to similar proteins^32^. Therefore, to investigate the binding environment (BE) of gliflozins, the resolved structures of the hSGLT2 (PDB entry: 7vsi^33^), protein-gliflozin and homologues of SGLT family or the same domain, sodium:symporter family (SSF, Pfam: PF00474^34^), are essential. However, its transmembrane characteristics raise difficulties in protein crystallography^35^. The multistage of transporters and the low similarity between homologues (sequence identity (id) < 30%)^36^ put challenges on the interaction analysis. Among the homologues with available structural data, *Vibrio parahaemolyticus* SGLT (vSGLT; PDB entry: 3dh4, 2xq2^37,38^) and hSGLT1 (PDB entry: 7sla, 7sla8^39^) share the same function of transporting pyranoses with hSGLT2. The pyranose BSs of these three proteins are highly resembling^33,39^. Yet their BSs accessibility for the solvent are completely opposite from that of the only resolved hSGLT2 (PDB entry: 7vsi^33^).

For the chemical structures of compound gliflozins, in our preliminary studies, similar compounds (Tanimoto coefficient > 0.85) show no targets other than SGLTs^40^. Furthermore, empagliflozin-hSGLT2 (PDB entry: 7vsi) is the single resolved structure in the databases today^41^, which decelerates a deeper understanding to the binding requirements for these compounds. Herein, we here propose a strategy to explore drug binding site (BS) under structural data shortage. Instead of directly exploring the BE of gliflozins, we dissected the structure of hSGLT2i compounds and identified the key BE of the decisive substructures. For further analysis, we developed a scoring function, propensity to distinguish the specific environment among the SGLT-alike proteins.

## Results

**Figure 1.**
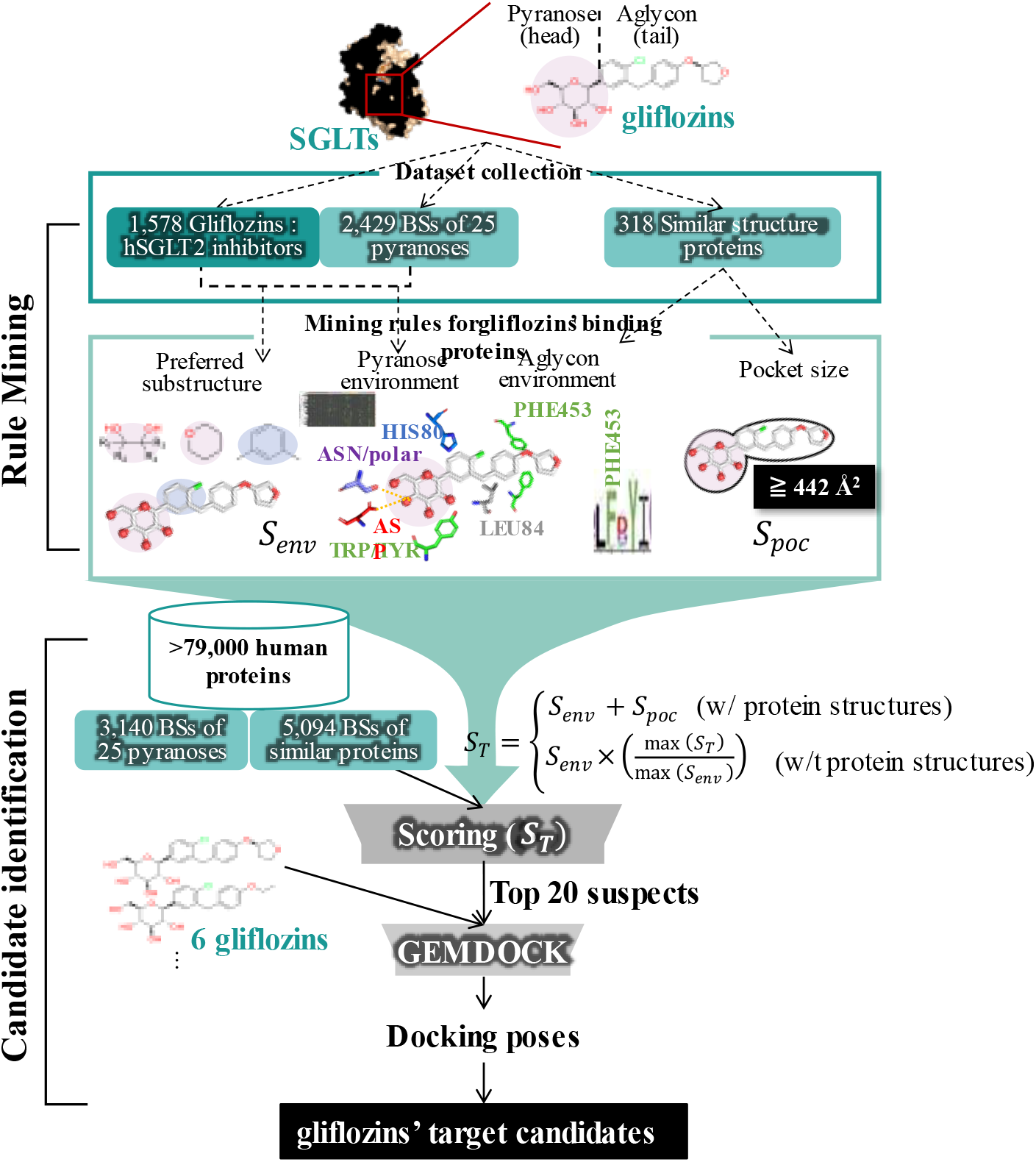
The overview pipeline. The figure demonstrates the key concepts and stops to identify the BE of gliflozins.

### Features of hSGLT2i compounds

We simply obtain two major features from the compounds of gliflozins. The number heavy atom (HA) and the key compound substructures. The size of the compound is highly related with its HA and can strongly affect cell permeability^42^ and its affinity to proteins. We examined the relationship between ligand HA, the conformation changes of proteins similar to hSGLT2, and the size of the protein pocket. These transporters transform from tunnels facing one side to the other at different stages^36,43^ to move compounds. The process of transition provides cavities of a variety of sizes. The conditions to accommodate ligands would change accordingly along the way^44^. We postulate that for binding pockets such as hSGLT2-empagliflozin, the larger the ligand, the larger the accommodating protein pocket is required. In the same type of proteins, there might be a reference interval of the pocket sizes for a given compound with a known number of HA, for example, stringent hSGLT2i in transmembrane transporters. To prove the hypothesis, we prepared the hSGLT2i grouped by different efficacy. The result demonstrated that the number of HA is statistically significant related to its efficacy in hSGLT2. This also indicates that compounds with efficacy concentrate >1000 nM (**Supplementary 1**) are distant from stringent hSGLT2i.

By comparing the hSGLT2i with non-hSGLT2i and the preference of each substructure to present in a hSGLT2i by odds-ratios, we were able to identify the key functional groups to form a hSGLT2i. **Figure 2**BC shows the top 10 substructures that determine a hSGLT2i, which largely corresponds to the approved gliflozins. Especially for the substructures highlighted in pink, they are the same with the 12 functional groups dissected from pyranoses (**Supplementary 2**). This result largely corresponds to the compound analysis on approved gliflozins (**Figure 2**A). Herein, this proved that pyranose is a critical part for the hSGLT2i and provided us a lead to explore the BE of gliflozins. By studying the common interactions between pyranoses and their binding proteins, we can obtain the key proportion of environment to stabilize a gliflozin. According to **Figure 2**C, it is possible to foresee that pyranose may lack of electrostatic force, but strong in van der Waals (vdW) force field (Ring in drug #006, i.e., tetrahydropyran) and hydrogen bond (H-bonds) provided by the hydroxyl groups.

**Figure 2.**
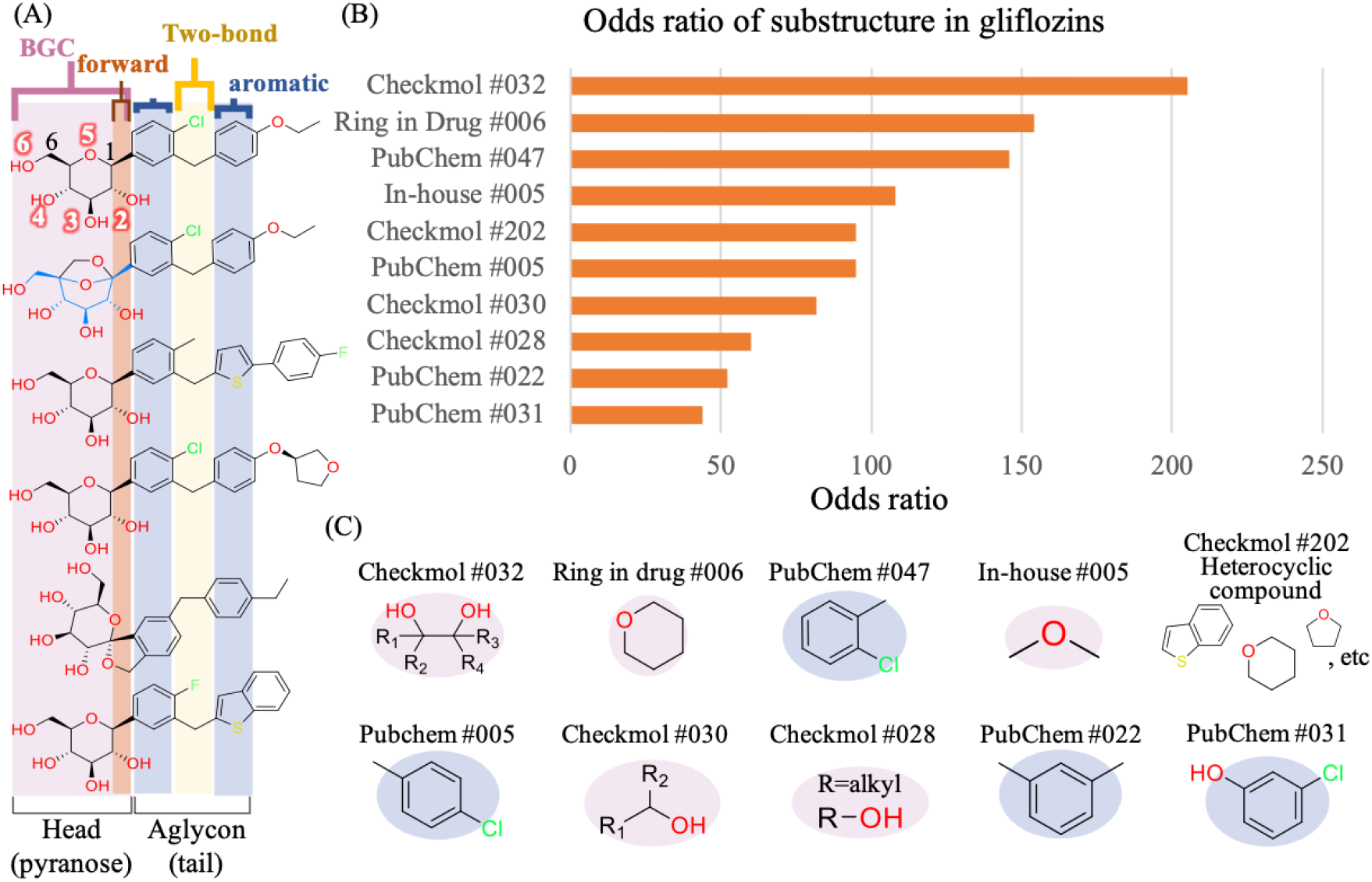
Important substructures in gliflozins. (A) Compound analysis on the approved gliflozins. β-D-glucopyranose (BGC) is also the original substrate of hSGLT2. The numbers are labelled based on the atom numbering of pyranoses (**Supplementary 3**). (B) The substructures with top ten odds ratios against non-hSGLT2i. (C) The substructures listed according to the odds ratios’ order. Substructures colored in pink overlap with those dissected from pyranoses, i.e., the BGC part in the gliflozins. The substructures highlighted in blue refers to the aromatic part in the gliflozins.

### The van der Waals BE of pyranoses

We collected 25 pyranoses from The Protein Data Bank (PDB^41^), 12 of them with resolved structures outside of glycosylation (**Supplementary 4**). Interestingly, the ligands that hSGLT2 and vSGLT transport take more than 80% of the 1,572 pyranose BSs, which are glucose and galactose. This refers to the normality of glucose and galactose BSs in pyranose-interacting protein environment and enables us to gather a large-scale dataset of gliflozin-relevant BS environment. The interaction profile in **Figure 3**A shows the interacting patterns of 1,572 pyranose BSs. Apart from the pyranose BSs, we put hSGLT2-empagliflozin as a reference to observe the pyranose binding environment. Note that the corresponding atom of pyranose oxygen 1 (O1) in empagliflozin is substituted with a carbon (C) (**Figure 2**A). The energy at the atom is calculated accordingly, i.e., it provides no H-bonds. vSGLT-GAL (PDB entry: 3dh4) is included in the 1,572 BSs and lies in the same cluster of hSGLT2-empagliflozin via 1-Pearson’s average correlation. This indicates the highly similar interaction patterns of the two BSs.

The interaction shows that the majority of the pyranose BSs participate the vdW linked with aromatic residues and H-bonds linked to negatively charged residues. The result matches with the roles of the negative residues at O4 and aromatic rings hydrophobic for pyranose binding. For instance, in *Neisseria* amylosucrase (PDB entry: 1jg9, not shown), D144 forms a salt bridge and additionally stabilizes the boundary of glucose. Meanwhile, F250 furnishes a stacking hydrophobic contact to glucose^45^. Additionally, we noticed there were equally-distant energy performance on some residues. (Yellow boxes in **Figure 3**) The same patterns are extremely strong on aromatic residues in multiple clusters prevalently, as we have mentioned the majority of BSs had vdW linked to these residues. Since the interaction is aromatic-residue-specific on pyranoses, we termed it “ring” stacking (**Figure 4**B). Multiple sequence alignment (MSA) had evinced that although hSGLT2 and vSGLT shared little sequence similarity as homologues, the interacting residues in pyranose BSs were still highly conserved. Based on MSA, the of SGLT homologues (**Figure 4**A), the consecutive (vSGLT) Y263 and W264 in pyranose BS are highly conserved across species. Next, we were curious if there were any preference of the pyranoses BSs to certain type of aromatic residues.

The multiplied energy (see Method) of the atoms on tetrahydropyran against each of the four aromatic residues are shown as **Figure 3**B. The higher HA of TRP is theoretically doomed to result in a higher performance on the index. However, when comparing the multiplied energy of TYR and that of PHE, we still observed a great deal of differences on both the count of pyranose BSs and the quantity of their performance on these two residues with close HA numbers (**Figure 3**C). These elaborate the pivotality of residues that are both polar and aromatic, TYR and TRP, in pyranose BSs and SGLTs, which is a critical rule for gliflozin BE. The tendency was verified by the propensity scoring (**Supplementary 5**). Each atom in the tetrahydropyran in pyranoses was preferred equivalently and strongly in by the TYR and TRP. The preference was specific for the pyranoses BSs.

**Figure 3.**
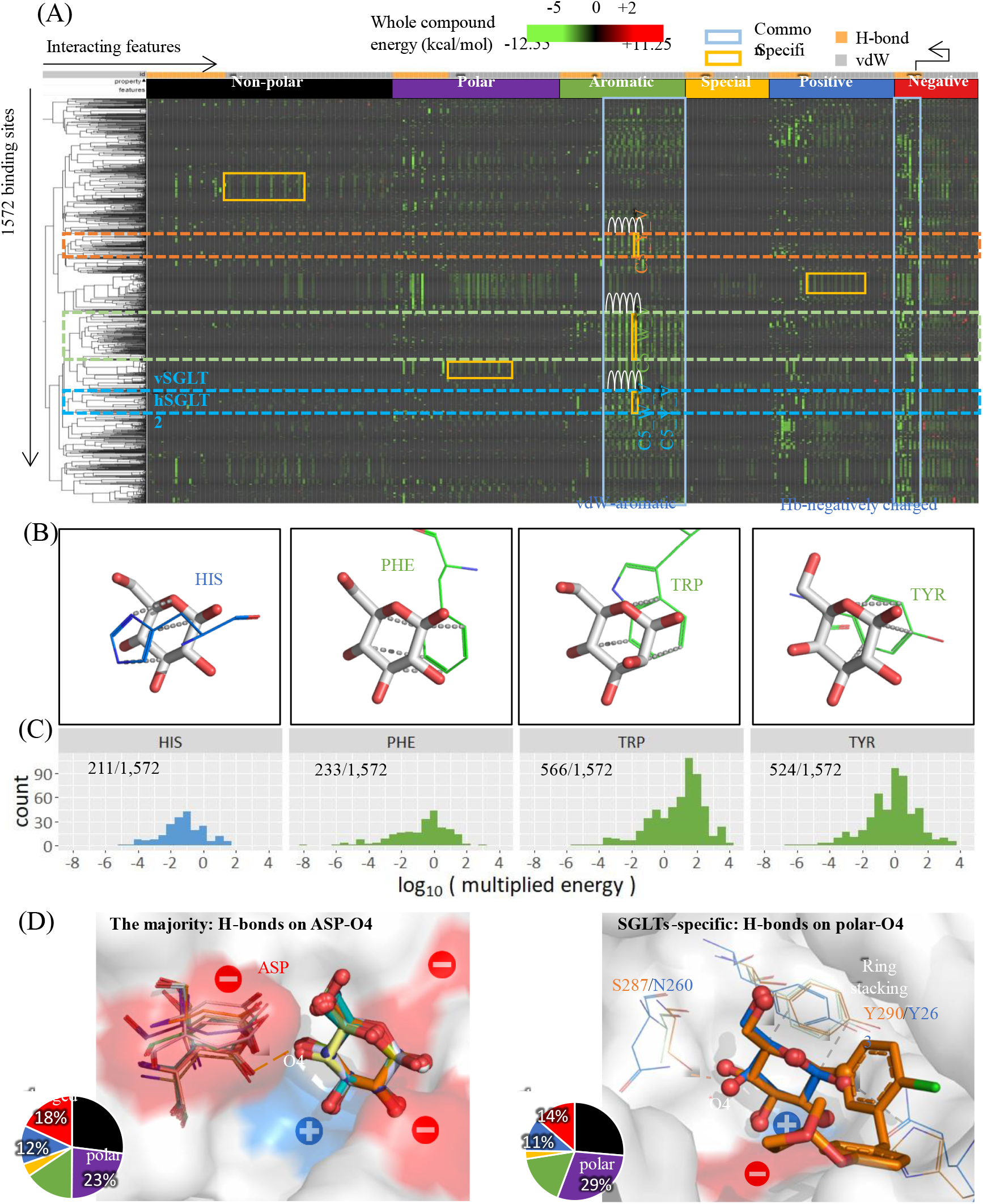
Pyranose interacting environments (A) Interaction profile of 1,572 pyranose binding environments (BEs). The first row indicates the force type: H-bonds (yellow-orange) or vdW (gray). The second row indicates the property of the interacting residue. A cell represents the assembled energy force in a binding site (see Material and Method). Partial compound in hSGLT2-empagliflozin complex was taken as a control in the analysis. The yellow boxes highlight the equally-distant energy on same residues in specific groups of pyranose BSs. The three clusters circled by dashed of colors point out the typical strong aromatic “ring” stacking field. The heat map is generated by morpheus GENE-E^46^. (B) Examples of BEs that demonstrate the major interaction patterns among the pyranose, which are aromatic residues providing “ring” stacking hydrophobic field. PDB entries from left to right: 3ft4, 2pc8, 6byg, and 3dh4. (C) The multiplied vdW force field of four aromatic residues to the tetrahydropyran substructure (multiplied energy, see Method) were drawn in grouped histograms. The box colors are mapped to the properties of the residues. (D) (Left) The majority (47.6%) of pyranoses BEs form H-bonds with ASP at O4. (Right)The specific group of BEs resembling hSGLT2. These BSs accommodate gliflozins and construct H-bonds at O4 with polar residues instead of ASP. These two groups show a statistically significant difference (X^2^: 93.791, p <2.2e-16) in their composition of BE residues. The charged residues are denoted by red(negative)/blue(positive) colored surfaces.

## The Hydrogen bonds of BE in pyranose BSs

For the H-bonds environment in pyranose BSs, we’ve seen the majority constructs H-bonds with negatively charged residues. Interestingly, though O1-O4 in pyranose seem to be in similar stereo-orientation, the preferences show that H-bonds frequent on O4 and secondly O3 (**Supplementary 6**). Among them, over 45% are contributed by ASP, which is denoted as O4_D_H. By comparing the phenomenon of this O4_D_H interaction, we identified that those of SGLTs-alike BEs were unique. On **Figure 3**D, one can see that the both pyranose-binding SGLTs render a polar residue to O4, instead of an ASP. The observation was supported by MSA. It indicates that it is crucial for SGLT-alike proteins to provide a polar force on the corresponding position (**Figure 4**A). According to Niu *et al*^*33*^., and Faham *et al*., ^38^, there are only a common pair of positive and negative charged residue in the pyranose BE to stabilize each other and the pyranose ligand.

To investigate deeper, we calculated the percentage of the residues of each property. The pie charts in **Figure 3**D demonstrate that solely the residue providing H-bond to O4 may be decisive for the charge-property of the entire pyranose BE. As in the majority group, these BSs form H-bonds to O4 with ASP, they have a much higher percentage of charged residues in their BSs, especially those negatively charged. For the SGLT-specific, known to accommodate gliflozins and providing polar residues to O4, show a frequency of polar residues in the entire BSs. In a previous mutagenesis study^39^ modifying H83Q (H80 in hSGLT2) on hSGLT1, the protein aborted function of sugar transport completely. Together with the fact that sporadic charged residues are extremely rare in SGLT-BSs, evidence indicates the subtle charged balance is crucial for the BEs for pyranose, and may further influence the affinity with gliflozin.

**Figure 4.**
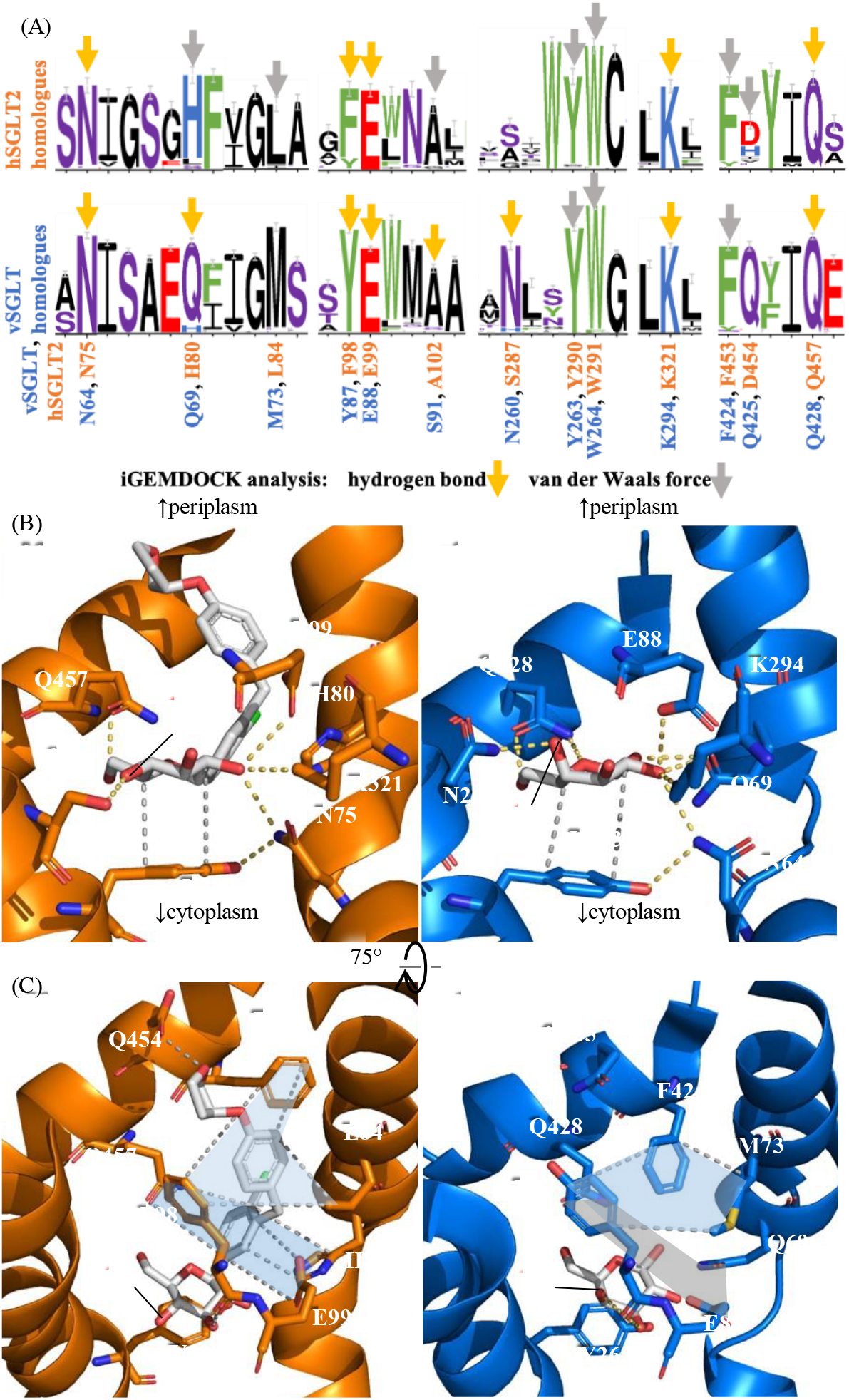
Key interacting residues influencing affinities of proteins to gliflozins. (A) The conservation analysis on the empagliflozin and GAL binding residues in hSGLT2 and vSGLT respectively. The residues are colored according to their properties. Based on the interaction algorithm of iGEMDOCK^47^, the residues providing H-bond environments are pointed with yellow-orange arrows; the residues form vdW field are pointed by grey arrows. (B) The binding residues stabilizing the pyranose or tetrahydropyran substructure in the two homologues work alike. (C) The key and corresponding residues interact with the aglycon part of empagliflozin. vSGLT is shown in marine blue cartoon/sticks; hSGLT2 is shown in orange cartoon/sticks. vdW force fields are labelled in grey dashes and filled with blue polygons; H-bonds are visualized in yellow-orange dashes. The grey polygon labelled on vSGLT-GAL indicates the missing hydrophobic field exists in hSGLT2-empagliflozins complex. Figures are visualized in PyMOL^48^.

### BE of hydrophobic field for the tail of gliflozins

According to **Figure 2**, we know that aromatic rings, especially 6-member rings, linked afterwards the pyranose substructure are critical to form an effective hSGLT2i. In hSGLT2, these aromatic rings are surrounded by strong hydrophobic field formed by non-polar or aromatic residues, especially for the second phenyl group^33^. Meanwhile, the first phenyl substructure is stabilized by H80, Q457, and E99. The latter two residues construct strong H-bonds with the pyranose substructure simultaneous at the orientation (**Figure 4**BC). In MSA (**Figure 4**A), we can see that these residues (H80, L84, F98, E99, F453, and Q457 on hSGLT2) are highly conserved in homologues of hSGLT2, which includes hSGLT1. Medical chemists have shown the inhibitory effects of gliflozins and phlorizin on hSGLT1, which are weaker than those to hSGLT2 but still efficacious (IC_50_ < 10^4^ nM)^49^, while phlorizin shows no inhibitory effects (IC_50_ > 10^6^ nM)^50^ to vSGLT. By juxtaposing the conservation analysis, we see the greater difference occurs on H80 (Q69 in vSGLT, **Figure 4**A). H80 stacks with the first phenyl group. It is the same substructure that substitutes O1 of β-glucose. Thus, we assumed that a BS provided a favorable environment for gliflozins if it had aromatic residues at the O1 position of pyranose or the corresponding residues of H80.

### Pocket size of gliflozins take at least 442 Å ^2^

In terms of homopharma, similar proteins provide similar pocket sizes for similar-sized compounds. Based on the same logic, beside from the BEs, we were able to predict the pocket size required to contain a gliflozin. As mentioned earlier, there were limitations on the homologues of hSGLT2 due to their low similarity in BSs. After the launch of the hSGTL2 structure, we had enough resource and turned to structural similarity. Among the 278 protein chains of known resemblance in structural biology, 180 contain a ligand in the center of the transmembrane proteins, just as GAL in vSGLT. They provide the materials to establish the relationship between ligands HA and protein pocket sizes. The pocket size of a BS is in strong flexibility to adapt their binding ligands to facilitate each process^51^, which is much determined by the protein functions and the conformation stages. Instead of the entire cavity constructed by the “proteins”, we are interested in the space to accommodate the “ligands”. Thus, we fetched 8Å-radius and 6Å -radius BSs in these similar proteins. When proteins are measured in whole chains, pocket sizes fluctuated under the same HA of ligands as predicted. We suspect that it is much serious because (1) the multistage and transitions of SSF and similar proteins, just as the open tunnel in inward vSGLT^38^ or (2) the original structural difference of different proteins. The pockets approximate the space of ligand occupation the best at 6Å -radius BSs and the ligands are well covered. We see the pocket sizes grow with increasing numbers of HA.

Further, we applied the desired HA interval of gliflozins. The evidence indicated the least space needed to fully hold a compound within the HA was 491Å ^2^. Taking the account that proteins are in large flexibility and change of movement, we set the cut-off line at 442 Å ^2^ (0.9 × 491) for the filter of off-target candidates.

**Figure 5.**
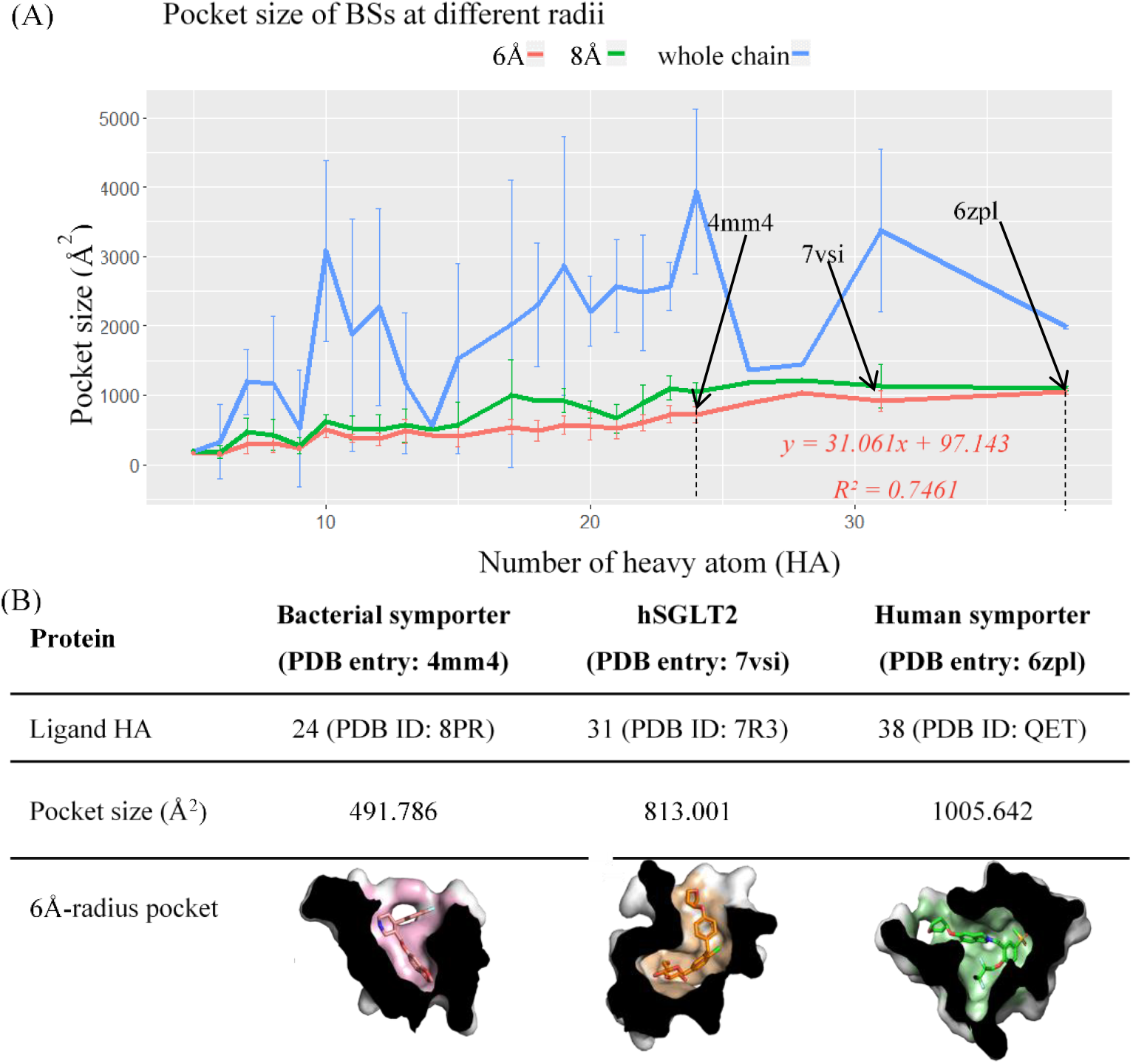
Pocket size mining for gliflozins. (A) According to the 180 hSGLT2-similar proteins, 6Å -BSs exhibit a linear relationship on pocket size with HA of ligand. The dashed lines indicate the HA interval of hSGLT2i. (B) The pocket sizes accommodating the ligands within the interval correspond to the linear relationship. As a ligand gets larger, it requires a larger pocket size in the protein. The last row illustrates the BSs in section views by PyMOL^48^.

### Identifying possible candidates

Based on the investigation above, we constructed a scoring function (*S*_*T*_) to assess the possibility of a BS for gliflozin. A number of > 7,900 human proteins were considered as pre-candidates. After further filtration, 8,237 BSs originated from 55 proteins were evaluated based on *S*_*T*_. To understand the functioning and the applicability of our investigation and the developed *S*_*T*_ we conducted a hit-rate analysis. The 10 proteins in the verifying set (details in Material and Method) were known to be inhibited by the stringent hSGLT2i (**Supplementary 1**) and taken as the positive set in **Figure 6**.

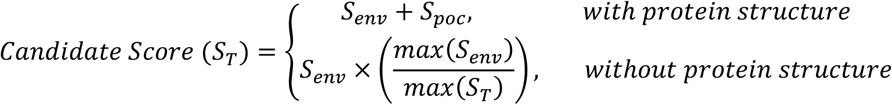

Our scoring function comprises the BE and the physical space of a BS to assess its capacity to bind a gliflozin. **Figure 6**BC indicate the known gliflozin BS (hSGLT2-empagliflozin) ranked high in the both criteria. We observe that the *S*_*T*_ scoring system yields better at predicting gliflozin binding pocket in **Figure 6**D. BE has always played a decisive role for ligands boundary. Our *S*_*env*_ has successfully captured the key BE of gliflozins as it does to the known protein target of gliflozins. Though *pocket size* may be a factor for a BS’s affinity to gliflozin, but both theoretically and the result demonstrate itself solely would not be adequate for gliflozin’s boundary to a BS. Based on the principle of bioinformatics, proteins with similar functions tend to share similar BE in pockets^31,52^. The prototype before *S*_*T*_ included the concept of functional similarity (**Figure 6**D, *Raw S*_*T*_ *+ S*_*func*_). On *Raw S*_*T*_ and *Raw S*_*T*_ *+ S*_*func*_, we see a similar pattern on curve of hit rate. Note that the BE, *S*_*env*_, takes the greater proportion in *S*_*T*_. Additionally, when *S*_*env*_ *+ S*_*func*_ is taken as the scoring function (not shown), it shows the same hit rate as *S*_*env*_. This indicates annotations on function similarity largely duplicate the BE of the candidates and it is not reasonable to rate it twice. To integrate the physical space of the pocket size into consideration but not to neglect the lack of resolved structures of the known BSs at the meanwhile, the final *S*_*T*_ was produced and it exceeds the others as shown in **Figure 6**D.

**Figure 6.**
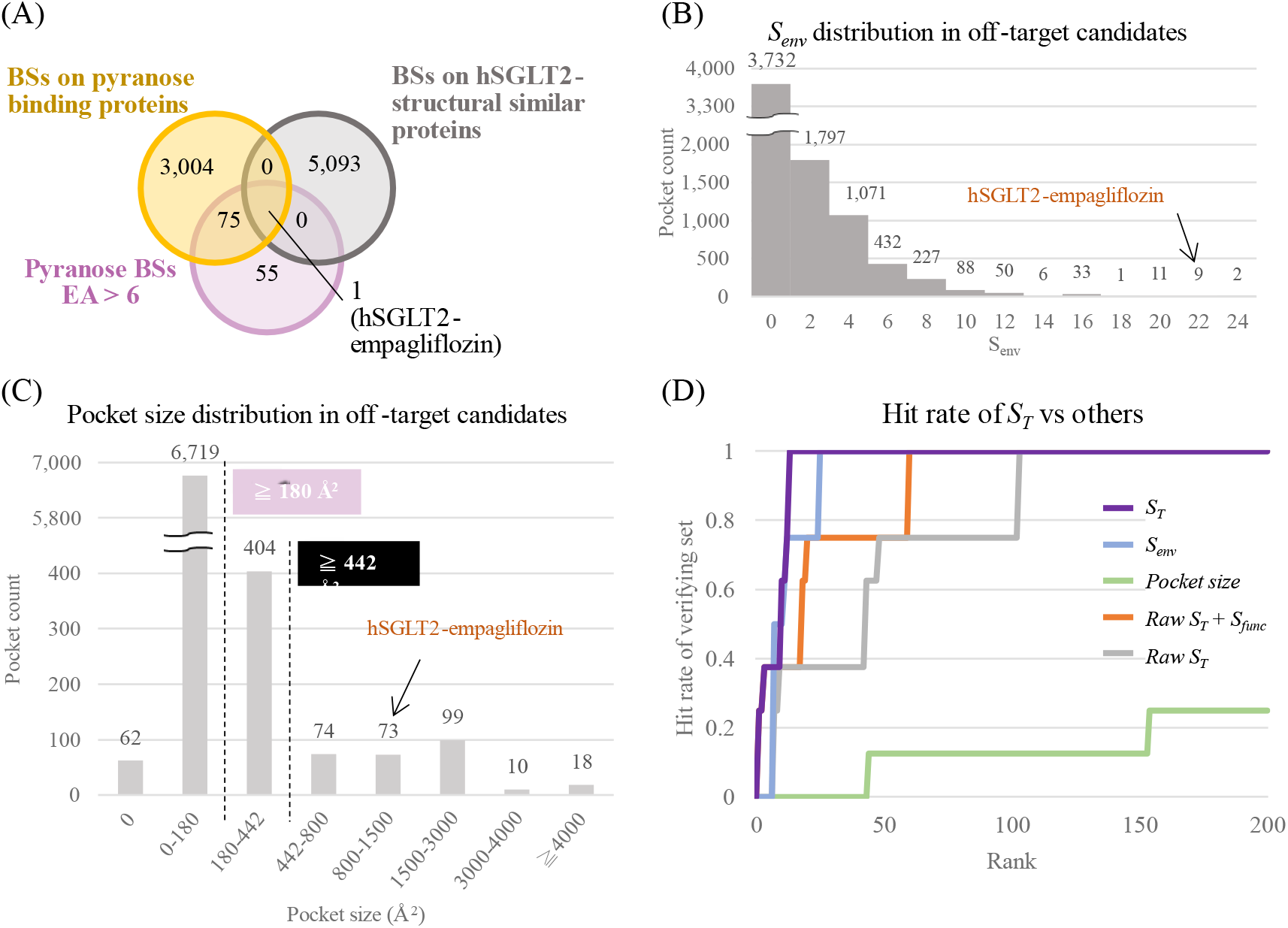
Performance of S_T_. (A) Composition of the pre-candidate set. The BSs are comprised by BSs from hSGLT2 structurally similar proteins and pyranose-binding proteins (see Material and Method). (B) The pocket size distribution of 8,237 pre-candidate BSs. The BSs with undetectable pockets are included in 0. (C) The S_env_ distribution of pre-candidate BSs. 180 Å ^2^ is the least pocket size required to hold a pyranose (see). (D) *Pocket size* indicates the direct volume of each BS. *Raw S*_*T*_ is the scoring function when a BS without accessible pocket size is not evaluate separately. *Raw S*_*T*_ *+ S*_*func*_ is the performance of a modified scoring system developed on *Raw S*_*T*_. *S*_*func*_ denotes the degree of how a BS share the same GO annotations in SGLTs, which is associated to its functional similarity to hSGLT2. *S*_*env*_ represents the term evaluating the BE solely in *S*_*T*_.

The BS with the best performance in each protein was further selected as the final candidates. We docked empagliflozin and five other common acknowledged approved gliflozins into the top-ranked BSs using iGEMDOCK^47^. The pose examples of empagliflozin were shown. An interesting candidate from pyranose BS outstands from the others (**Figure 7**B). The protein participates in the cardiovascular system protection. CBG is the abbreviation for cytosolic β-glucosidase. Back in the 1990s, researchers had known the beneficial effects of grape juice and wine on CVD were due to the compounds called trans-piceid and trans-resveratrol^53,54^. In 2006, Henry-Vitrac *et al*. demonstrated that the organelle with CBG hydrolyzed trans-piceid and produced trans-resveratrol. The same effect was found on the plasma membrane extract with lactase phlorizin hydrolase (LPH)^55^. We suspected that gliflozins were hydrolyzed by CBG/LPH and underwent the same pathways that are beneficial for CVD.

To prove this hypothesis, it is essential for the gliflozin to fit into the active site of CBG. The strong hydrophobic field providing residues (W345, Y309, F225, and V168) were mentioned to stablize the glucoside proportion as resveratrol of a glucose derivatives^56^. The same residues resemble the enironment formed by H80, L87, F98, and F453 in hSGLT2 and accommadate docked empagliflozins. They attract the benzene parts of gliflozins. Both negatively charged (E424/E373) and polar (Q17) residues furnish H-bonds to BGC and empagliflozin on O4. W417 constructs a “ring” stacking environment for the pyranose substructure, as we have discussed in the last chapter. Especially for E373 and W417, the two residues have been proposed to be critical for the active site of CBG^56^. The observation correlates with the rules concluded in this investigation and offers gliflozins a potential BS.

Another example comes from a structural similar protein situated in renal tubule membrane, sodium-dependent neutral amino acid transporter (B^0^AT1). The protein is responsible for the amino acid transport in intestine and kidney^57^. Especially in the renal system, it functions similarly with SGLT2 to reabsorb substance from the urine. The docked pose (**Figure 7**D) indicates that the two benezene substructures of gliflozin can be stabilzed by R57, a residue that is charged but also with a higher HA to provide adequate vdW force. The mutation on R57 is closely associated with Hartnup disorder^58^ and thus possibly at the BS for transporting amino acids. The patient is known to produce urine with undegraded amino acids^57^, just as individuals with mutation on SGLT2 have presistent glucosuria^3^. The docked pose and the previous research together illustrate that it is possible that gliflozins bind to the tranport BS of B^0^AT1 and block the protein like how they inhibit SGLT2.

**Figure 7.**
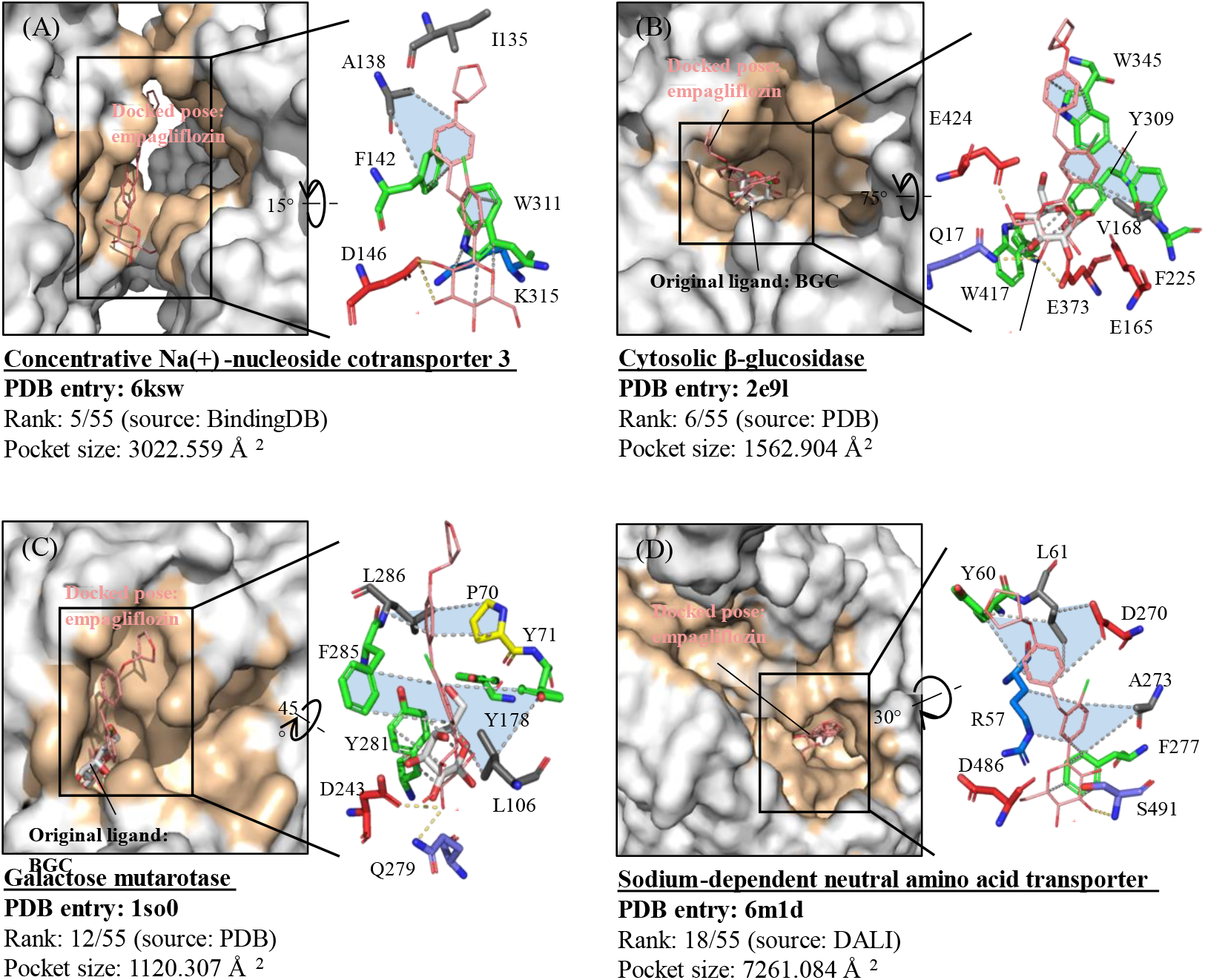
Docked poses show the possibility of gliflozins’ boundary in proteins. (A) A protein known to interact with gliflozins. (B) The BE of docked empagliflozin in Cytosolic-β-Glucosidase (PDB entry: 2e9l) active BS and key binding residues. The wheat surface refers to the predicted binding pocket, which is 1562 Å 2. (C) Another pyranose BS shown potential to bind with gliflozins. (D) A renal protein sharing structural similarity with hSGLT2. The compound shown in white/red sticks is the original glucose in the resolved structure. The salmon lines demonstrate the docked pose of empagliflozin. The yellow orange dashed lines visualized the possible H-bonds, while blue areas lined with gray dashes indicate vdW force field. The four section views are drawn in difference scales. The figures are displayed with PyMOL^48^.

## Conclusions

So far, we have found the physical condition of a protein pocket ≧ 442 Å ^2^ to enable the occupation of gliflozins by applying similar transmembrane transporters. For the analysis of gliflozins’ BE, we have overcome the obstacles of data shortage in resolved structures by identifying the key substructures. The common pyranose-substructure provided us with > 1500 BSs, for further BSs analysis. To explore the specific interactions in pyranose BSs and evaluate the decisive features for the gliflozin boundary, we developed a propensity score. Integrated with the result of the interaction profile analysis, we were able to recognize three types of binding patterns:

(1) The vdW “ring” stacking between aromatic residues and pyranose ring, particularly to TYR and TRP
(2) H-bonds on O4 of pyranose
  a. To ASP: the BS may consist more negatively charged residues
  b. To polar residues, especially ASN: a bigger proportion of the BS residues are polar and less proportion are negatively charged.

Despite of the data shortage, taking hSGLT2-empagliflozin as template and the combination of MSA, we were enabled to propose the importance of the hydrophobic field for the benzene substructures, particularly the one substituting O1 in pyranose. The further proof can be amplified in the research afterwards.

For the verification and application of these criteria, we have constructed a scoring function, *S*_*T*_ to evaluate the possible BSs with known binding proteins. Some of the top-ranked candidates are related with CVD or the normal functioning of kidney. Combined with the docked pose analysis, these BSs may explain the uncover mechanism of hSGLT2i. With the integration of gliflozins genomics data in the future, we are looking forward to finding the off-targets and uncovering the common mechanisms for the three chronic diseases. In general, the proposed methodology would help overcome the limitation of data shortage in predicting drug off-targets. Specifically, for gliflozin and its related diseases, the result can provide improved knowledge for the drug mechanisms.

## Material and Method

### Dataset collection: compound sets and their features

The compounds listed in **Figure 2**A are the gliflozins approved of treatment for DM in multiple countries. Though there are some other hSGLT2i been approved to treat DM. These six compounds are documented in at least two of the following: European Medicines Agency (EMA)^59^, U.S. Food and Drug Administration (U.S. FDA)^60^, Japan Pharmaceuticals and Medical Devices Agency (Pmda)^61^, and KEGG database^62^. They should be approved of treating type 2 DM as the standalone compound. If the gliflozins is approved as a part of a combination treatment, it will not be listed as one of the six.

Compound features were applied to provide mainly two aspects in this study: (1) the substructures required for a hSGLT2 inhibitor and (2) the number of heavy atoms required for an effective hSGLT2 inhibitor. Both were collected from bindingDB (2021m9)^40^ and filtered according to the conditions listed as below. The conditions finally resulted in 1,578 compounds, of which 1393 were contributed by records with IC_50_. Trimming with the following criteria, we resulted in 692 hSGLT2i.

(1) Targeted protein with uniprot ID: P31639^63^.
(2) Removal of duplicated compound was judged based on the replication of bindingDB monomer ID. Each record was kept in the following priority:
  a. The data with half maximum inhibitory concentration (IC_50_) > concentration 50% of the maximal effect (EC_50_) > inhibitory constant (K_i_).
  b. The most effectivity/ the least concentration record
(3) Data for K_i_ were removed for the limited information to provide for efficacy. The data of IC_50_ or EC_50_ were considered the same as “efficacy concentration (conc.)”.
(4) Data with none published record were removed for the less traceability and less informative.
(5) According to the IC_50_ of approved gliflozins^64,65^, we defined the efficacy conc. ≤ 1000 nM as hSGLT2i. The rest of the gliflozins with efficacy conc. >1000 nM to hSGLT2 were termed “hSGLT2 binding compounds”. They were presented as a set of negative reference for the HA distribution of hSGLT2i.

To understand the functional groups that affect the affinity of a compound for hSGLT2, we also prepared a negative set. We explored 990 commercial-available, random, drug-like compounds prepared by Yang *et al*.,^66^ and gathered 718 compounds shared the similar HA interval with stringent hSGLT2i (**Supplementary 1**). The HA interval of these non-hSGLT2i was set 20-42 for the tolerance of addition or substitution of a six-member ring. The numbers of negative compound set (718) and positive hSGLT2i compound set (692) are weighted closely, it thus requires no sampling to avoid imbalanced bias. Considering the lower performances of each individual system of molecular descriptors^67^, we utilized the integrated 553 descriptors according to Huang *et al*.^67^ to represent the substructures of gliflozins: (1) 204 fingerprints from checkmol^68^ (2) 168 PubChem fingerprint^69^ (3) 34 functional groups in drugs^70^ (4) 147 types of rings frequently seen in FDA drugs^71^. The important substructures were determined by their odds ratios. We counted the numbers of compounds with and without each substructure in both sets. Below shows a canonical example for 553 substructures. The process resulted in 174 substructures with meaningful odds ratios, since *b, c, or d* could be zero.

| Set | Positive set (692 hSGLT2i) |  | Negative set (718 non-hSGLT2i) |  |
| --- | --- | --- | --- | --- |
| Condition | with | without | with | without |
| Substructure # | *a* | *b* | *c* | *d* |

### Sequence sets and multiple sequence alignment (MSA)

Multiple sequence alignment is a critical technique for identifying the important residues of given proteins. If a fragment or a residue is crucial for the protein to maintain its key functions, we can anticipate that the same residue is kept during evolution and that high conservation of the fragments can be seen in homologues across species^72^. In the structures currently available with pyranose ligands, vSGLT is the closest homologue. In this study, the identification of critical residues in hSGLTs and vSGLT would provide information on the essential binding requirements of SGLT for inhibitors. Even a residue of the hSGLT2 counterpart in vSGLT is different, the combination of variance on two structures and MSA across other species would render the conservation and key residues to stabilize the boundary of pyranose or gliflozins.

The id between hSGLT2 and vSGLT is quite low as homologues (32%, BLASTP^73^) and the two organisms are distantly related, which may refer to the conservation difference in bacterial and eukaryotic SGLTs. For this reason, we gathered the sequences separately and conducted MSA independently. In this case, we would not have to worry if the sequences are biased for prokaryotes or eukaryotes. Additionally, for the balance of two sets, we took a nonredundant protein sequences dataset (NR, July 2021) for MSA analysis.

After BLASTP against NR, 5000 sequences were collected for in hSGLT2-driven and vSGLT-driven sequence set with E-value < 10^-137^. The coverages were > 51% and > 39%, and id were > 43% and 41%, respectively. Further trimming conditions are listed:

(1) To avoid those nonhomologous proteins of SGLT are included and hinder the conservation:
  a. sequence length coverage > 75%.
  b. Several samples of “unclear species”, replication of the same proteins, “predicted”, “hypothetical” or labeled “low qualities” were recorded. These sequences are removed.
  c. To avoid less informative sequences and sequences of unknown sources, sequences with description containing “MULTISPECIES”, unspecified “bacterium”, “candidate”, and “uncultured” are removed. These sequences were found mainly in vSGLT sets, where most of the samples are bacterial.
(2) To avoid mistaken conservation of replicated sequences, the following criteria were applied:
  a. 633 groups of eukaryotic isoforms were skimmed and one was retained in each group. Reasonably, these proteins were only found in hSGLT2 sets.
  b. The representative sequences of each cluster at 80% were selected after being trimmed to reduce computational load. (See below)

Finally, a sequence was selected for each species and each genus in the hSGLT2 and vSGLT sets, respectively, to reduce computational load. To reduce the id% bias and sequence replication in conservation analysis, a clustering of sequences was performed on CD-hit^74^. We tested the cut-off point for cluster id at 90% and 80%. We employed the latter, considering the yet high similarity of the sequences at the cutoff point of 80%. Sequences of hSGLT2 (uniprot ID: P31639, GenPept NCBI reference sequence NP_003032.1) and vSGLT (uniprot ID: P96169, GenPept NCBI reference sequence: WP_029825056.1) were checked to be included in the final sets. This resulted in 89 in hSGLT2-driven set and 253 sequences in vSGLT-driven set. They were further undergone MSA by T-COFFEE^75^.

Few bacterial sequences similar to SGLT were reviewed in Swissprot. Hence, the validation of two sets was performed separately, based on different databases. The validation of hSGLT2-driven set was conducted directly based on 32 sequences from Swissprot retrieved by BLASTP. The sequences were aligned by T-COFFEE. For the vSGLT-driven set, the entire uniprot database (Swissprot, TrEMBL and Protein Informative Resource) was taken into account, which was not an option available on BLASTP. Regarding the selection of the database, sequence selection and alignment of the vSGLT set was thus performed in ConSurf ^76^.

### Dataset collection: pyranose-co-crystalized structures, set pyr

According to the outcome of important features of hSGLT2 inhibitors, pyranoses were part of the critical functional groups to interact with hSGLT2. To extend the list and find the most potential candidates, we screened over 180,000 PDB structures (2021 August) to construct pyranose-co-crystallized protein structures. Pyranoses in chain glycans, glycosyl groups, or covalently bound with other substances were removed. Structures labeled with standalone pyranose were included. The rest were filtered by the type of “CONECT” record in the pdb files for covalent bonding to the pyranoses. For glycosylation, the complexes were excluded based on the structure description from PDB RCSB. The trimming criteria were:

(1) The ligand formula with “C 6 H 12 O 6” (C_6_H_12_O_6_) and the compound structure looked as **Supplementary 7**D.
(2) Completely co-crystalized pyranose with 12 atoms. Incomplete resolved pyranoses were removed.
(3) EA > 6. On the basis of previous unpublished research, our lab set up a scoring function to identify the ligand in the cavity.

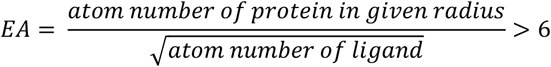

EA value evaluates the depth of the ligand by calculating the modified ratio in the given distance. Here, we set the radius at 4.5 Å. The deeper a ligand is located inside its protein, the greater the probability that it is enveloped by residues. With the same number of denominators, 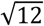, the numerator is thus larger.

These steps kept 1,572 pyranose BSs originating from 1,271 protein chains, 418 uniprot IDs. This structural set was termed “set pyr” and used further for mainly three purposes: (1) the large-scale of structural set provided as a large-scale data to analyze the interacting features of each specific pyranose-protein pair; (3) in order to explicate the functions and preference of pyranose-protein pair in the contrast with other proteins, this set also provided materials to establish a propensity of pyranose-atom to each specific type of amino acid.

### Dataset collection: protein similar to hSGLT2, set ps

Due to the lack of protein structures, molecular pathway annotations, and few known targets for gliflozins, we first focused on the proteins that share a high similarity with hSGLT2 to explore off-targets for gliflozins. The resolved homologues in bound forms shared little id beside from the vSGLT-GAL structure. After the launch of the hSGTL2 structure, we were able to better understand the tunnel binding environment of hSGLT2-similar proteins. We therefore turned to structural similarity conducted by DALI^77^ to find the proteins containing similar BS as hSGLT2 to stabilize gliflozins. We selected “PDB search” for collection of all possible similar against proteins structures in PDB. We received 364 protein chains in this step, which included all the so-far-known crystalized proteins of SSF family. Since only a few SSF proteins showed id > 30% to hSGLT2, id would not be a proper filter for further judgement of valid the result of structural similarity. To remove unwanted protein structures showing less consistency with hSGLT2, we enumerated simple criteria as below. The trimming process produced 280 chains of protein structures, composing set ps (protein similarity). Those with ligands (180 chains) provided resources for pocket size approximation on gliflozin compounds.

(1) Z-score – rmsd < 0, where rmsd stands for root-mean-square deviation of atomic positions to hSGLT2 and Z-score evaluates the similarity with the query structure, hSGLT2. The higher the Z-score is or the smaller the rmsd is, the more similar is the protein to the structure of hSGLT2.
(2) Alignment length/sequence length (seq ratio) > 0.5 We want the Z-score and rmsd calculated based on a fair proportion of the structure sequence length. The alignment length against hSGLT2 is required to be at least half of the sequence length of the given structure. When the ratio < 0.5, we may observe additional segments of protein unable to align with the other protein.
(3) Proteins must be well-studied enough to be recorded in uniprot, so further comparison with genomic data is feasible.

### Dataset collection: the pre-candidate of gliflozin off-target

To assess the capacity of BE affinity to gliflozins and for the interpretability of binding mechanism, we focused our targets on BS-basis. As discussed previously, the conformational variation would largely affect a BS to accommodate a ligand. Hence, resolved structures sharing a common Uniprot ID would be considered separately as a pre-candidate. Among the 7,900 human proteins, we concentrated on the hSGLT2-similar and pyranose-binding proteins to increase the hit and reduce computational load. There were 131 BSs originated from 126 human proteins within the 1,573 BSs in set pyr. For the 280 protein chains in set ps, 67 were human. These proteins were then processed by CASTp for the evaluation of each possible BS as a potential gliflozins BS and preliminary generated more than 34,172 BSs. However, not all of them qualified for further assessment.

It was critical that we have reference to assess the BE in these BSs. For the BSs other than those of set pyr, we employed paired-sequence alignment (T-coffee^75^) and mapping of corresponding residues on non-pyranose-BSs. Proteins in set ps acquired via structural similarity were applied accordingly by DALI structural alignment^77^. For instance, we’ve found it essential for gliflozins to anchor on H80, S287, W289, Y290, and W291 in hSGLT2. Based on sequence alignment, human ficolin-2 (Uniprot ID: Q15485) has residues^78^ of LYS, SER, TRP, HIS, and at the corresponding positions. Though the algorithm leaves W291 (hSGLT2) without a matching residue in Q15485, this largely helps us to evaluate the BE when it lacked of a ligand of pyranose. For a BS in ficolin-2 deep enough to be detected as a pocket, but without at least one of the residues above, the BS was to eliminate from the list due to the obstacles in BE assessment. If a protein performs extremely poorly aligned to hSGLT2 that no residue is mapped at the correspondence, all BSs from the protein were to delist as well.

### Interaction profile at pyranose-atom-based level

According to Lennard-Jones potential^79^, people today know that the extreme of vdW attraction is at about 6 Å and the minimum of H-bond is at about 2.3Å. It requires at least a 6Å radii to construct a BE. For the 1,572 BS in set pyr, we extracted each bound pyranose and the residues within 10Å-radii. Ligands and their bound BSs were isolated directly by the edition on text in pdb files. In **Supplementary 8**D, we see the consistency of 12 atoms in the SGLTs despite their stereoisomerisms. The aligned residues are still conserved in properties. The uniform orientation indicates the specificity of each atom in pyranose. In light of the diversity of the pyranose-binding proteins, we designed the features in the combination of three factors to unify the interacting characteristics (**Supplementary 9**), which are 12 pyranose atoms: C1-C6, O1-O6, 20 essential amino acids, and 3 major binding forces: electrostatic force (E), hydrogen bond (H, or H-bonds), van der Waals force (V, or vdW). With the composition of pyranose, the profile for the analysis of the binding environment contains 6 × 20 (H-bond) + 12 × 20 (vdW) = 360 features in the interaction profile generated by GEMDOCK algorithm^66^. The performance of a BS in each feature was denoted as an ensemble of the BS environment. If C1 in a pyranose furnishes vdW to multiple TYR, the interactions of all these vdW forces are summed up and rendered as one strong annotation in C1_W_V.

#### The multiplied energy to aromatic residues

As **Figure 3**AB had shown, the interaction profile displayed a pattern of vdW simultaneously happening to C1-C5, and O5 in a number of pyranose BSs. The six atoms construct the functional group of tetrahydropyran, which is “Ring in drug #006” in **Figure 2**C. This quickly links us to π-π stacking between aromatic rings^80^. Pyranose and the aromatic residues may furnish a resembling “ring” stacking hydrophobic interaction to all the atom on functional group of tetrahydropyran in these BSs. We simply evaluated this by multiplying all the energy on these pyranose atoms predicted by GEMDOCK^66^ in pyranose to aromatic residues shown as the formula (HIS, PHE, TRP, and TYR).

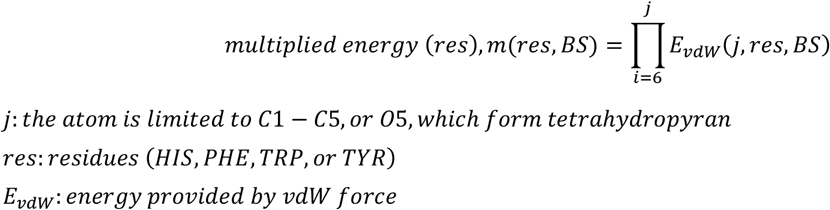

Let’s suppose the output of a given pyranose in its BS, *pyr1 in bs*, has a multiplied vdW energy to TYR of *m*. Via the multiplication, if there is a paucity of vdW force in any of the six atoms of *pyr1*, it can be easily identified at *m=0*. If TYR forms a (or odds number of) repulsive boundary to one of the six atoms in *pyr1, m < 0*. When *bs* forms a ring stacking hydrophobic environment with TYR or when *bs* consists of several TYR surrounding the ligand at a proper distance, *m > 0*. In this case, *bs* provides an environment resembling SGLTs, the binding proteins of gliflozins. The performances were weighted based on the HA of the aromatic residues and their preference in SGLTs BS. The weighted performances of the four aromatic residues were further put into a summation to evaluate the stacking environment of the BS.

#### The distribution of the properties of the residues in BSs

In the analysis of O4-Hb, the two groups of BSs were isolated from set pyr. The majority of the BSs laid in the group of O4-ASP. This included any pyranose BS that form a H-bond interaction with ASP at energy < 0 (kcal/mol). The SGLT-alike group was comprised of the BSs developing H-bonds to any polar residue rather than any ASP at energy < 0 (kcal/mol). The two groups were composed of 749 and 226 BSs, respectively. Based on Lennard-Jones potential, it is deducible that the smallest radius to construct a complete BS for a ligand would be 6 Å . As a result, we calculated the residues with any atom inside the 6Å -radii BS in set pyr. The property of a residue was classified according to that listed in **Supplementary 10**.

### Propensity of pyranose to protein-binding sites

Although hSGLT2 and vSGLT transport different types of pyranoses, the alignment demonstrated the consistent orientation of the sugars (**Figure *4***BC). We thus hypothesized that each O in pyranose may be similar as subgroups, e.g., hydroxyl, but unique in its stereo-orientation, or subtle chemical property. By establishing pyranose propensity analysis, we are able to evaluate the preference of each O and C in the pyranoses to each amino acid. To prevent statistical biases caused by similar proteins, it is required to identify a group of representing pyranoses within set pyr. In set pyr, 1,572 BSs were further studied in the interaction and were hierarchically clustered by average 1-Pearson correlation. BSs were aligned by Pymol^81^ for further comparison. The result was manually observed for similarity of the interaction and the interaction profiles to determine 379 clusters. Each cluster generated a representative BS for further analysis.

(1) In singleton clusters, the BS was identified as a representative.
(2) For the others, each BS would be calculated for its interacting similarity with the others in the same cluster by Pearson’s correlation. The one with the highest average similarity in each cluster was chosen as representatives.

A universal protein set as the normalizing background is needed. We will call it set b. Set b provided surface/BS residues to stand for any protein other than pyranose binding ones, by which could help obtain the specificity of each interaction within pyranose BSs. To explore the specificities of the propensity of these interactions in pyranose binding environment, we modified 30%^82-85^ thinning set of ProteinNet 7 established by *AlQuraishi* ^86^ and took it as the representative set of the other random proteins (set b). Structures of obsolete PDB entries were removed. The structures of pyranose-binding proteins listed in set pyr were excluded from set b. The structures of overlapping sequences were removed to avoid static bias. The 30% thinning was further verified by CD-hit^74^ at 30% and provided 6,284 clusters. A representative structure was selected for each cluster. After a series of trimmings, set b comprised 6,284 protein chains. We utilized NACCESS^87^ to measure the exposure of each residue and its access from the protein surface. The parameters of a surface residue were regulated in the “absolute” accessibility (acc.) of the “1.4Å”-radius-solvent at “2.5Å^2^”. The side chain count on GLY in set b was omitted due to its composition as a single hydrogen (**Supplementary 11**).

The construction of the propensity in this study was inspired by the formalism of BLOSUM62^88^, DrugScore^PDB84^, and PCA-Pred^82^. We followed the equation of Gibb’s free energy for a system at chemical equilibrium^89^ to assess the preference of a pyranose for its binding protein^84^:

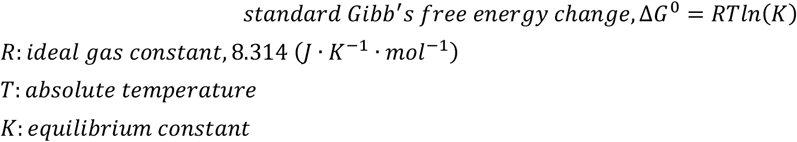

In our approach, we simplified the idea and defined the propensity of pyranose atoms to amino acids in BS as follows:

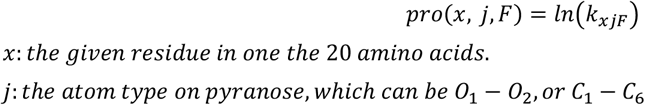

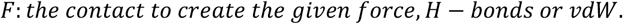

*k*_*xjF*_ was the main part annotating the specificity of the amino acid preference in pyranose BSs. We have:

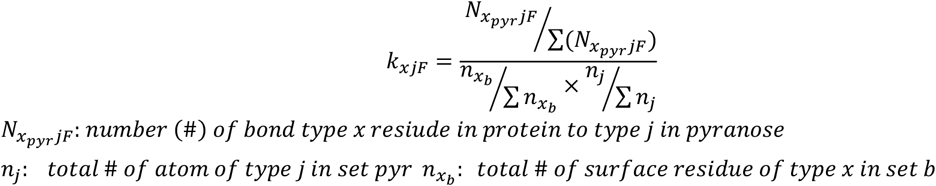

By applying set b, we were able to create a normalized ratio of certain amino acids in pyranose BSs and on other proteins surfaces, *i*.*e*., possible binding residues for ligands. Since the preference for a given type of atom in pyranose in BS was individually quantified, we considered it crucial to balance it in the number of HA. This would develop a constant of 1/12, *n*_*j*_/∑ *n*_*j*_.

Let’s assume we are interested in quantifying the specificity of the H-bonds between GLU in proteins and O1 in pyranoses, as exhibited in **Supplementary 12**B. We calculate the H-bonds of O1-GLU furnished in each pyranose BS. We repeat the procedure on 379 representative pyranose BSs to obtain 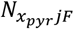. The calculation was carried out repeatedly on any other *x* residue in proteins and *j* atom in pyranoses and produced 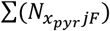. Finally, we acquired *pro(O1, Q, H)*. Upon normalization with the terms in **Supplementary 13**A-1 and **Supplementary 14**A-2 and taken under natural logarithm, the propensity demonstrated the specificity of H-bonds on O1-GLU in pyransoe compared to other protein BSs.

### Pocket size and pocket mapping

#### The approaches to measuring a pocket size

There are two well-used algorithms for measuring the size of the protein pocket. Connolly’s molecular surface area, or solvent-excluded surface (SES). The area is defined as the surface on which the solvent molecule of radius R is in contact with the van der Waals surface of a given protein.^90^ Richard’s surface, or solvent accessible surface area (SAS). The surface consists of the points where the centers of the solvent molecules of radius R are placed during the movement on a given protein. Theoretically, majority part of the solvent is water (H_2_O) and R is 1.4 Å at default^91^. The existence of a ligand in a given space will influence the distribution of any compounds in the same space. Since the distributions of water molecules are flexible in the pocket, we consider SES more satisfactory to the pocket measurement of gliflozins.

We compared two widely used pocket size measuring tools: fpocket^92^ and CASTp^93^. Both calculate pocket sizes by the occupation of alpha spheres^94^, which is based on alpha-shape theory, first developed in 1994. The former provides pocket size according to SES, while the latter provides both SAS and SES. fpocket is available on the command line terminal and is more straightforward for large-scale analysis of BSs. There are also few differences between the analyses of whole protein or divided individual chains. However, some situations were observed (**Supplementary 15**): (1) the distances of the spheres that make up the same pocket were too far in some tunnels (2) the fpocket does not appear to support the DALI output files. (3) The weak adaptivity of fpocket in narrow tunnels/channels

CASTp developed an unfavorable outcome when whole protein is taken into account. The same situation was observed in other known pocket size measuring tools, for instance, SURFNET^95^, a pocket measurement tool available on PDBsum^41^. This was somehow inevitable due to the optimized algorithm and restrictions in biological crystallization to produce homomeric or heteromeric protein chains that may not exist naturally. We were able to overcome the problem simply by parsing the content of the pdb files into divided chains. The measurement on CASTp was then chosen in this study after an overall assessment.

#### Predicting the pocket sizes of similar-sized compounds

We postulated that for pockets as deep as hSGLT2i in the tunnel and surrounded by protein residues, compounds of similar size will occupy similar pocket sizes in proteins. As the compound HA grows, the size of the pocket may expand. In terms of homopharma, similar proteins provide similar pocket sizes for similar-sized compounds. The HA interval within mean (μ) ± 2 × standard deviation (σ) of set stringent hSGLT2i was applied as the HA for compounds similar to gliflozins (**Supplementary 1**). The 180 protein structures with ligands in the set ps contributed to the similar protein and BSs. The ligands were required to be embedded enough in protein BSs to form detectable pockets. This was evaluated by EA > 6 and CASTp. We demanded that their ligand HA annotation on PDB should be the same as their resolved crystal structure to be informative for pocket size prediction. The 180 ligand BSs were enough for us to build and prove the linear relation between a HA of ligand and its spined pocket size.

When measuring the whole pocket size to accommodate GAL in vSGLT-GAL structure, we found that the entire tunnel of a transporter was likely to be recognized as a pocket. These compounds obviously did not require the pocket size of an entire tunnel to form a stable boundary with proteins. The pockets on off-targets for gliflozins may not necessarily be a tunnel either. The largest distance to form a vdW connection is 6 Å . The average length of a main chain of an amino acid is approximately 3.5Å ^96^. To be more preservative, we thus shortened the length to 2Å as the gap and tested the predicted pockets starting from 6 Å, 8 Å, and 10 Å (**Figure 5**).

### Selection for gliflozin off-target candidates

Here we explained the composition of the scoring function to find the gliflozin off-target BS. The pre-candidate set was majorly composed by the BSs from human proteins in set pyr and set ps. Owing to the fact that we evaluated the BE through unidentical method, it becomes essential to integrate the two. In brief, our scoring function can be denoted by a simple equation:

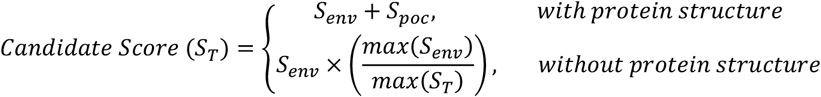

#### Rating for protein pocket sizes

In two origins of the pre-candidates, the pocket sizes were measured with the same principle. For the resolved structures without ligand, we were not able to draw the corresponding pocket size of the BS to form the interactions in current method. They were rated as 0. We had 491 Å ^2^ for the least volume to hold the compound with similar HA to the smallest hSGLT2i in hSGLT2 similar proteins. The threshold was determined at 491 × 0.9 = 442 to adapt the possible movement of pockets. As a result, we had:

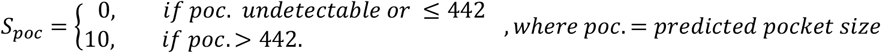

#### Interactions on key residues

The term of *S*_*env*_ comprised the score for ring stacking environment (S_*stacking*_), the score to evaluate O4-H-bonds (*S*_*O4-H*_), and the score rating π-π stacking to the benzene substructures at O1 substitute (*S*_*O1-vdW*_). The three subterms were assessed separately in set pyr and set ps.

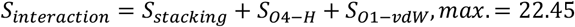

### Set pyr

For the ring stacking between aromatic residue and tetrahydropyran substructure on pyranose, we concentrated on TYR to hSGLT2 (**Figure 3**). A BS energy performance obtained from the pyranose interaction profile has a lot of information to provide about its BE. However, it is necessary to standardize the information with the BE prediction in set ps. Since *m*(*TYR*, 7*vsi*_*A*_7*R*3) = 17.3 and the reduction on dimensionality was required, the cut-off for a conspicuous ring stacking was set at > 15. Additionally, taking into account (1) the present of each aromatic residue in set pyr, (2) the interaction in SGLTs, (3) the HA associated with the contribution to vdW(**Supplementary 16**), the weights in the scoring function were set accordingly.

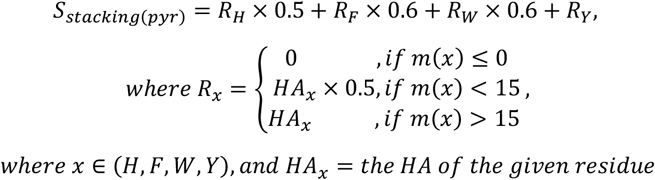

The H-bonds environments on O4 and hydrophobic environment for the benzene in gliflozins were included in the scoring function. In the O4-H environment, we’ve seen that polar residue is a preferred H-bond partner to gliflozins-specific BE, while ASP frequent the most in all pyranose BSs.

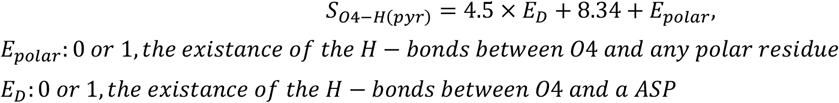

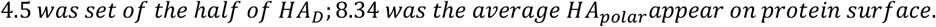

On the other hand, considering that H80 (hSGLT2) stabilized the functional group substituting O1 in empagliflozin (**Supplementary 17**) in a manner of π-π stacking, aromatic residues were favored at O1 vdW environments. F98 and Y290 also provide a strong hydrophobic field to the substituting benzene. We thus defined the threshold at > 2 aromatic residues and had:

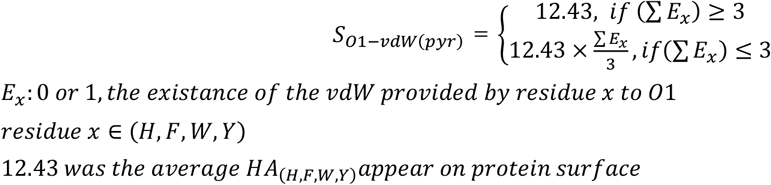

### Set ps

Since different ligands in the structures share protein similarity with hSGLT2, the interactions of these proteins for gliflozins environments were evaluated based on the structural alignments instead. For the “ring” stacking subterm (*S*_*stacking*_), the residues (*R*) aligned to W289 (*R*_*W289*_), Y290, and W291 were rated. For H-bonds on O4, properties of the residue aligned with S287 were assessed for H-bond preference; for the π-π stacking with the benzene substructures in gliflozins, residues corresponding to H80 (hSGLT2) were weighted. Again, the weights were adjusted according to the importance of each feature for gliflozins boundary and the balance of scores between each term and subterm. As a result, we have:

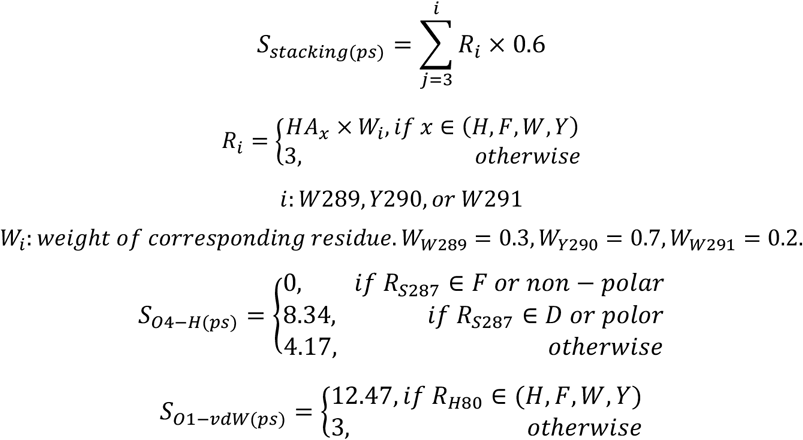

The same transformation to *S*_*T*_ performed on set ps was applied on the BSs with BE evaluated by sequence alignment. The scores of set pyr and set ps were then able to ranked together despites of their different predicting approach. In both transformation of scoring function, hSGLT2 (7vsi) was ranked as top 1 with the score of 39.55.

#### Composition of verifying set

Previously, we’ve collected 692 hSGLT2i for BindingDB^40^. In order to build up a positive set of BSs that is known to interaction with hSGLT2, we reversed the searching filters and looked for the proteins with experimental data on these 692 gliflozins. We acquired 1,392 records on 22 unique proteins across five species, which are *Homo sapiens* (human), *Mus musculus* (mouse), *Rattus norvegicus* (rat), *Sus scrofa* (pig), *Agaricus bisporus* (white button mushroom). Again, we merged records on IC_50_ and EC_50_ to produce efficacy conc. as we did to <u>Dataset collection: compound sets and their features</u>. The datum with the least efficacy conc. on each protein was annotated as its effectivity. Protein with efficacy conc. < 20 μM were included as the positive controls.

It is worth mentioning that in this study, we are measuring the possibility of each BS rather than a protein to bind with gliflozins. Since the BS on these proteins that hold gliflozins are still unclear, the *S*_*poc*_ for them are ranked by the largest pocket detected on each protein. Protein without resolved structure had 0 on *S*_*poc*_. Regarding that none of these proteins overlapped with the proteins in set pyr, the BE of these BS were evaluated based on the sequence alignment as we did in <u>Dataset collection: the pre-candidate of gliflozin off-target</u>. The composition of the pocket residues was examined manually to be included in the aligned residue to H80, S287, W289, Y290, and W291 on hSGLT2.

### Virtual screening with gliflozin compounds

The compounds used to represent gliflozins have been described in <u>Dataset collection: compound sets and their features</u>. In **Figure 7**, only empagliflozin was shown, considering the hSGLT2-empagliflozin structure is available as a comparative reference. The structures of these compounds were acquired from the isomeric SMILE in PubChem^97^ for manual analysis (**Figure 2**A). The drugs were transformed to 3D format for docking preparation by Open Babel^98^. Protein BSs are extracted with Swiss-PDBViewer^99^ manually taking the pocket detected by CASTp^93^ as template and 10 Å -radii as the main principle. The docking condition was set uniformly with 100 poses from 70 generations among 1,000 of population.

## Reference

1 Estimates, G. H. The top 10 causes of death (2000-2019). (WHO, 2020).

2 Harding, J. L., Pavkov, M. E., Magliano, D. J., Shaw, J. E. & Gregg, E. W. Global trends in diabetes complications: a review of current evidence. Diabetologia 62, 3–16, doi:10.1007/s00125-018-4711-2 (2019).

3 Vallon, V. The mechanisms and therapeutic potential of SGLT2 inhibitors in diabetes mellitus. Annu Rev Med 66, 255–270, doi:10.1146/annurev-med-051013-110046 (2015).

4 Grundy, S. M. Metabolic syndrome: connecting and reconciling cardiovascular and diabetes worlds. J Am Coll Cardiol 47, 1093–1100, doi:10.1016/j.jacc.2005.11.046 (2006).

5 Prevention, C. f. D. C. a. Estimates of Diabetes and Its Burden in the United States, <https://www.cdc.gov/diabetes/data/statistics-report/index.html> (2022).

6 Alicic, R. Z., Rooney, M. T. & Tuttle, K. R. Diabetic Kidney Disease: Challenges, Progress, and Possibilities. Clin J Am Soc Nephrol 12, 2032–2045, doi:10.2215/CJN.11491116 (2017).

7 Lovshin, J. A. et al. Retinopathy and RAAS Activation: Results From the Canadian Study of Longevity in Type 1 Diabetes. Diabetes Care 42, 273–280, doi:10.2337/dc18-1809 (2019).

8 Emerging Risk Factors, C. et al. Diabetes mellitus, fasting blood glucose concentration, and risk of vascular disease: a collaborative meta-analysis of 102 prospective studies. Lancet 375, 2215–2222, doi:10.1016/S0140-6736(10)60484-9 (2010).

9 (U.S. Food and Drug Administration, 2020).

10 (U.S. Food and Drug Administration, 2021).

11 EMA/642015/2021 (European Medicines Agency, 2012).

12 Washburn, W. N. Development of the renal glucose reabsorption inhibitors: a new mechanism for the pharmacotherapy of diabetes mellitus type 2. J Med Chem 52, 1785–1794, doi:10.1021/jm8013019 (2009).

13 Ehrenkranz, J. R., Lewis, N. G., Kahn, C. R. & Roth, J. Phlorizin: a review. Diabetes Metab Res Rev 21, 31–38, doi:10.1002/dmrr.532 (2005).

14 Lee, J. et al. Novel C-aryl glucoside SGLT2 inhibitors as potential antidiabetic agents: 1,3,4-Thiadiazolylmethylphenyl glucoside congeners. Bioorg Med Chem 18, 2178–2194, doi:10.1016/j.bmc.2010.01.073 (2010).

15 Hummel, C. S. et al. Glucose transport by human renal Na+/D-glucose cotransporters SGLT1 and SGLT2. Am J Physiol Cell Physiol 300, C14–21, doi:10.1152/ajpcell.00388.2010 (2011).

16 Scirica, B. M. et al. Saxagliptin and cardiovascular outcomes in patients with type 2 diabetes mellitus. N Engl J Med 369, 1317–1326, doi:10.1056/NEJMoa1307684 (2013).

17 Zinman, B. et al. Empagliflozin, Cardiovascular Outcomes, and Mortality in Type 2 Diabetes. N Engl J Med 373, 2117–2128, doi:10.1056/NEJMoa1504720 (2015).

18 Perkovic, V. et al. Canagliflozin and Renal Outcomes in Type 2 Diabetes and Nephropathy. N Engl J Med 380, 2295–2306, doi:10.1056/NEJMoa1811744 (2019).

19 Heerspink, H. J. L. et al. Dapagliflozin in Patients with Chronic Kidney Disease. N Engl J Med 383, 1436–1446, doi:10.1056/NEJMoa2024816 (2020).

20 Bhatt, D. L. et al. Sotagliflozin in Patients with Diabetes and Chronic Kidney Disease. N Engl J Med 384, 129–139, doi:10.1056/NEJMoa2030186 (2021).

21 Heerspink, H. J., Perkins, B. A., Fitchett, D. H., Husain, M. & Cherney, D. Z. Sodium Glucose Cotransporter 2 Inhibitors in the Treatment of Diabetes Mellitus: Cardiovascular and Kidney Effects, Potential Mechanisms, and Clinical Applications. Circulation 134, 752–772, doi:10.1161/CIRCULATIONAHA.116.021887 (2016).

22 Staels, B. Cardiovascular Protection by Sodium Glucose Cotransporter 2 Inhibitors: Potential Mechanisms. Am J Med 130, S30–S39, doi:10.1016/j.amjmed.2017.04.009 (2017).

23 Lytvyn, Y., Bjornstad, P., Udell, J. A., Lovshin, J. A. & Cherney, D. Z. I. Sodium Glucose Cotransporter-2 Inhibition in Heart Failure: Potential Mechanisms, Clinical Applications, and Summary of Clinical Trials. Circulation 136, 1643–1658, doi:10.1161/CIRCULATIONAHA.117.030012 (2017).

24 Action to Control Cardiovascular Risk in Diabetes Study, G. et al. Effects of intensive glucose lowering in type 2 diabetes. N Engl J Med 358, 2545–2559, doi:10.1056/NEJMoa0802743 (2008).

25 Duckworth, W. et al. Glucose control and vascular complications in veterans with type 2 diabetes. N Engl J Med 360, 129–139, doi:10.1056/NEJMoa0808431 (2009).

26 Zoungas, S. et al. Severe hypoglycemia and risks of vascular events and death. N Engl J Med 363, 1410–1418, doi:10.1056/NEJMoa1003795 (2010).

27 Rosenstock, J. et al. Cardiovascular outcome trials in type 2 diabetes and the sulphonylurea controversy: rationale for the active-comparator CAROLINA trial. Diab Vasc Dis Res 10, 289–301, doi:10.1177/1479164112475102 (2013).

28 Diaz-Rodriguez, E. et al. Effects of dapagliflozin on human epicardial adipose tissue: modulation of insulin resistance, inflammatory chemokine production, and differentiation ability. Cardiovasc Res 114, 336–346, doi:10.1093/cvr/cvx186 (2018).

29 Sato, T. et al. The effect of dapagliflozin treatment on epicardial adipose tissue volume. Cardiovasc Diabetol 17, 6, doi:10.1186/s12933-017-0658-8 (2018).

30 (U.S. Food and Drug Administration, 2022).

31 Chiu, Y. Y. et al. Homopharma: A new concept for exploring the molecular binding mechanisms and drug repurposing. Bmc Genomics 15, doi:10.1186/1471-2164-15-S9-S8 (2014).

32 Agamah, F. E. et al. Computational/in silico methods in drug target and lead prediction. Brief Bioinform 21, 1663–1675, doi:10.1093/bib/bbz103 (2020).

33 Niu, Y. et al. Structural basis of inhibition of the human SGLT2-MAP17 glucose transporter. Nature 601, 280–284, doi:10.1038/s41586-021-04212-9 (2022).

34 Mistry, J. et al. Pfam: The protein families database in 2021. Nucleic Acids Res 49, D412–D419, doi:10.1093/nar/gkaa913 (2021).

35 Ostermeier, C. & Michel, H. Crystallization of membrane proteins. Current Opinion in Structural Biology 7, 697-701, doi:10.1016/S0959-440X(97)80080-2 (1997).

36 Wahlgren, W. Y. et al. Substrate-bound outward-open structure of a Na(+)-coupled sialic acid symporter reveals a new Na(+) site. Nat Commun 9, 1753, doi:10.1038/s41467-018-04045-7 (2018).

37 Watanabe, A. et al. The mechanism of sodium and substrate release from the binding pocket of vSGLT. Nature 468, 988–991, doi:10.1038/nature09580 (2010).

38 Faham, S. et al. The crystal structure of a sodium galactose transporter reveals mechanistic insights into Na+/sugar symport. Science 321, 810–814, doi:10.1126/science.1160406 (2008).

39 Han, L. et al. Structure and mechanism of the SGLT family of glucose transporters. Nature 601, 274–279, doi:10.1038/s41586-021-04211-w (2022).

40 Liu, T., Lin, Y., Wen, X., Jorissen, R. N. & Gilson, M. K. BindingDB: a web-accessible database of experimentally determined protein-ligand binding affinities. Nucleic Acids Res 35, D198–201, doi:10.1093/nar/gkl999 (2007).

41 Berman, H. M. et al. The Protein Data Bank. Nucleic Acids Research 28, 235–242, doi:10.1093/nar/28.1.235 (2000).

42 Matsson, P. & Kihlberg, J. How Big Is Too Big for Cell Permeability? Journal of Medicinal Chemistry 60, 1662–1664, doi:10.1021/acs.jmedchem.7b00237 (2017).

43 Weyand, S. et al. Structure and molecular mechanism of a nucleobase-cation-symport-1 family transporter. Science 322, 709–713, doi:10.1126/science.1164440 (2008).

44 Coleman, J. A., Green, E. M. & Gouaux, E. X-ray structures and mechanism of the human serotonin transporter. Nature 532, 334–339, doi:10.1038/nature17629 (2016).

45 Mirza, O. et al. Crystal structures of amylosucrase from Neisseria polysaccharea in complex with D-glucose and the active site mutant Glu328Gln in complex with the natural substrate sucrose. Biochemistry 40, 9032–9039, doi:10.1021/bi010706l (2001).

46 Morpheus. <https://software.broadinstitute.org/morpheus> (

47 Hsu, K. C., Chen, Y. F., Lin, S. R. & Yang, J. M. iGEMDOCK: a graphical environment of enhancing GEMDOCK using pharmacological interactions and post-screening analysis. BMC Bioinformatics 12 Suppl 1, S33, doi:10.1186/1471-2105-12-S1-S33 (2011).

48 Schrodinger, L. The PyMOL Molecular Graphics System, Version 1.8 (2015).

49 Abdul-Ghani, M. A., DeFronzo, R. A. & Norton, L. Novel hypothesis to explain why SGLT2 inhibitors inhibit only 30-50% of filtered glucose load in humans. Diabetes 62, 3324–3328, doi:10.2337/db13-0604 (2013).

50 Xie, Z., Turk, E. & Wright, E. M. Characterization of the Vibrio parahaemolyticus Na+/Glucose cotransporter. A bacterial member of the sodium/glucose transporter (SGLT) family. J Biol Chem 275, 25959–25964, doi:10.1074/jbc.M002687200 (2000).

51 Stank, A., Kokh, D. B., Fuller, J. C. & Wade, R. C. Protein Binding Pocket Dynamics. Accounts of Chemical Research 49, 809–815, doi:10.1021/acs.accounts.5b00516 (2016).

52 Bass, J. I. F. et al. Using networks to measure similarity between genes: association index selection. Nature Methods 10, 1169–1176, doi:10.1038/nmeth.2728 (2013).

53 Pace-Asciak, C. R., Hahn, S., Diamandis, E. P., Soleas, G. & Goldberg, D. M. The red wine phenolics trans-resveratrol and quercetin block human platelet aggregation and eicosanoid synthesis: Implications for protection against coronary heart disease. Clinica Chimica Acta 235, 207-219, doi:10.1016/0009-8981(95)06045-1 (1995).

54 Romero-Pérez, A. I., Ibern-Gómez, M., Lamuela-Raventós, R. M. & de la Torre-Boronat, M.C. Piceid, the Major Resveratrol Derivative in Grape Juices. Journal of Agricultural and Food Chemistry 47, 1533–1536, doi:10.1021/jf981024g (1999).

55 Henry-Vitrac, C., Desmouliere, A., Girard, D., Merillon, J. M. & Krisa, S. Transport, deglycosylation, and metabolism of trans-piceid by small intestinal epithelial cells. Eur J Nutr 45, 376–382, doi:10.1007/s00394-006-0609-8 (2006).

56 Berrin, J.-G. et al. Substrate (aglycone) specificity of human cytosolic beta-glucosidase. Biochemical Journal 373, 41–48 (2003).

57 Milne, M. D. Hartnup disease. Biochem J 111, 3p–4p, doi:10.1042/bj1110003p (1969).

58 Kleta, R. et al. Mutations in SLC6A19, encoding B0AT1, cause Hartnup disorder. Nature Genetics 36, 999–1002, doi:10.1038/ng1405 (2004).

59 EMA/265224/2016 (European Medicines Agency, 2016).

60 (ed U.S. Food and Drug Administration) (2015).

61 (ed Pharmaceuticals and Medical Devices Agency) (Japan, 2019).

62 Kanehisa, M. & Goto, S. KEGG: kyoto encyclopedia of genes and genomes. Nucleic Acids Res 28, 27–30, doi:10.1093/nar/28.1.27 (2000).

63 UniProt, C. UniProt: the universal protein knowledgebase in 2021. Nucleic Acids Res 49, D480–D489, doi:10.1093/nar/gkaa1100 (2021).

64 Abdul-Ghani, M. A., DeFronzo, R. A. & Norton, L. Novel Hypothesis to Explain Why SGLT2 Inhibitors Inhibit Only 30–50% of Filtered Glucose Load in Humans. Diabetes 62, 3324–3328, doi:10.2337/db13-0604 (2013).

65 Mascitti, V. et al. Discovery of a clinical candidate from the structurally unique dioxabicyclo[3.2.1]octane class of sodium-dependent glucose cotransporter 2 inhibitors. J Med Chem 54, 2952–2960, doi:10.1021/jm200049r (2011).

66 Yang, J. M. & Chen, C. C. GEMDOCK: a generic evolutionary method for molecular docking. Proteins 55, 288–304, doi:10.1002/prot.20035 (2004).

67 Huang, Y. W. et al. Discovery of moiety preference by Shapley value in protein kinase family using random forest models. BMC Bioinformatics 23, 130, doi:10.1186/s12859-022-04663-5 (2022).

68 Haider, N. Functionality pattern matching as an efficient complementary structure/reaction search tool: an open-source approach. Molecules 15, 5079–5092, doi:10.3390/molecules15085079 (2010).

69 PubChem. PubChem Substructure Fingerprint. (2009).

70 Hsu, Y.-C. Moiety-based Site-moiety Map Master thesis, National Chiao Tung University, (2011).

71 Taylor, R. D., MacCoss, M. & Lawson, A. D. Rings in drugs. J Med Chem 57, 5845–5859, doi:10.1021/jm4017625 (2014).

72 Capra, J. A. & Singh, M. Predicting functionally important residues from sequence conservation. Bioinformatics 23, 1875–1882, doi:10.1093/bioinformatics/btm270 (2007).

73 Altschul, S. F., Gish, W., Miller, W., Myers, E. W. & Lipman, D. J. Basic local alignment search tool. J Mol Biol 215, 403–410, doi:10.1016/S0022-2836(05)80360-2 (1990).

74 Huang, Y., Niu, B., Gao, Y., Fu, L. & Li, W. CD-HIT Suite: a web server for clustering and comparing biological sequences. Bioinformatics 26, 680–682, doi:10.1093/bioinformatics/btq003 (2010).

75 Di Tommaso, P. et al. T-Coffee: a web server for the multiple sequence alignment of protein and RNA sequences using structural information and homology extension. Nucleic Acids Res 39, W13–17, doi:10.1093/nar/gkr245 (2011).

76 Ashkenazy, H. et al. ConSurf 2016: an improved methodology to estimate and visualize evolutionary conservation in macromolecules. Nucleic Acids Res 44, W344–350, doi:10.1093/nar/gkw408 (2016).

77 Holm, L. Using Dali for Protein Structure Comparison. Methods Mol Biol 2112, 29–42, doi:10.1007/978-1-0716-0270-6_3 (2020).

78 Endo, Y., Sato, Y., Matsushita, M. & Fujita, T. Cloning and characterization of the human lectin P35 gene and its related gene. Genomics 36, 515–521, doi:10.1006/geno.1996.0497 (1996).

79 Lennard-Jones, J. E. Cohesion. Proceedings of the Physical Society 43, 461–482, doi:10.1088/0959-5309/43/5/301 (1931).

80 Hunter, C. A. & Sanders, J. K. The nature of. pi.-. pi. interactions. Journal of the American Chemical Society 112, 5525–5534 (1990).

81 Schrodinger, LLC. The AxPyMOL Molecular Graphics Plugin for Microsoft PowerPoint, Version 1.8 (2015).

82 Siva Shanmugam, N. R., Jino Blessy, J., Veluraja, K. & Gromiha, M. M. Prediction of protein-carbohydrate complex binding affinity using structural features. Brief Bioinform 22, doi:10.1093/bib/bbaa319 (2021).

83 Taroni, C., Jones, S. & Thornton, J. M. Analysis and prediction of carbohydrate binding sites. Protein Eng 13, 89–98, doi:10.1093/protein/13.2.89 (2000).

84 Gohlke, H., Hendlich, M. & Klebe, G. Knowledge-based scoring function to predict protein-ligand interactions. J Mol Biol 295, 337–356, doi:10.1006/jmbi.1999.3371 (2000).

85 Malik, A. & Ahmad, S. Sequence and structural features of carbohydrate binding in proteins and assessment of predictability using a neural network. BMC Struct Biol 7, 1, doi:10.1186/1472-6807-7-1 (2007).

86 AlQuraishi, M. ProteinNet: a standardized data set for machine learning of protein structure. BMC Bioinformatics 20, 311, doi:10.1186/s12859-019-2932-0 (2019).

87 Hubbard, S. & Thornton, J. NACCESS (The University of Manchester, 1992).

88 Henikoff, S. & Henikoff, J. G. Amino acid substitution matrices from protein blocks. Proc Natl Acad Sci U S A 89, 10915–10919, doi:10.1073/pnas.89.22.10915 (1992).

89 Gautam, R. & Seider, W. D. Computation of Phase and Chemical-Equilibrium .1. Local and Constrained Minima in Gibbs Free-Energy. Aiche Journal 25, 991–999, doi:DOI 10.1002/aic.690250610 (1979).

90 Richards, F. M. & Fm, R. Areas, volumes, packing, and protein structure. (1977).

91 D’Arrigo, J. S. Screening of membrane surface charges by divalent cations: an atomic representation. Am J Physiol 235, C109–117, doi:10.1152/ajpcell.1978.235.3.C109 (1978).

92 Le Guilloux, V., Schmidtke, P. & Tuffery, P. Fpocket: an open source platform for ligand pocket detection. BMC Bioinformatics 10, 168, doi:10.1186/1471-2105-10-168 (2009).

93 Tian, W., Chen, C., Lei, X., Zhao, J. & Liang, J. CASTp 3.0: computed atlas of surface topography of proteins. Nucleic Acids Res 46, W363–W367, doi:10.1093/nar/gky473 (2018).

94 Edelsbrunner, H. & Mucke, E. P. 3-Dimensional Alpha-Shapes. Acm T Graphic 13, 43–72, doi:Doi 10.1145/174462.156635 (1994).

95 Laskowski, R. A. SURFNET: a program for visualizing molecular surfaces, cavities, and intermolecular interactions. J Mol Graph 13, 323-330, 307-328, doi:10.1016/0263-7855(95)00073-9 (1995).

96 Ainavarapu, S. R. K. et al. Contour Length and Refolding Rate of a Small Protein Controlled by Engineered Disulfide Bonds. Biophys J 92, 225-233, doi:10.1529/biophysj.106.091561 (2007).

97 Kim, S. et al. PubChem in 2021: new data content and improved web interfaces. Nucleic Acids Res 49, D1388–D1395, doi:10.1093/nar/gkaa971 (2021).

98 O’Boyle, N. M. et al. Open Babel: An open chemical toolbox. Journal of Cheminformatics 3, 33, doi:10.1186/1758-2946-3-33 (2011).

99 Guex, N. & Peitsch, M. C. SWISS-MODEL and the Swiss-PdbViewer: an environment for comparative protein modeling. Electrophoresis 18, 2714–2723, doi:10.1002/elps.1150181505 (1997).

