## Supplementary Data for "Off-target prediction of SGLT2 inhibitors: an integrative bioinformatics approach to uncover structural mechanisms"

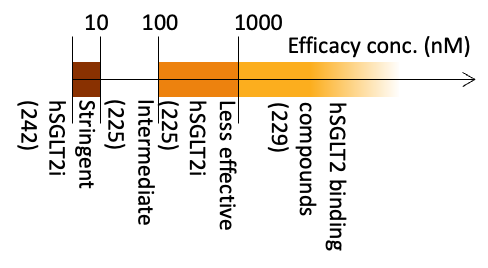

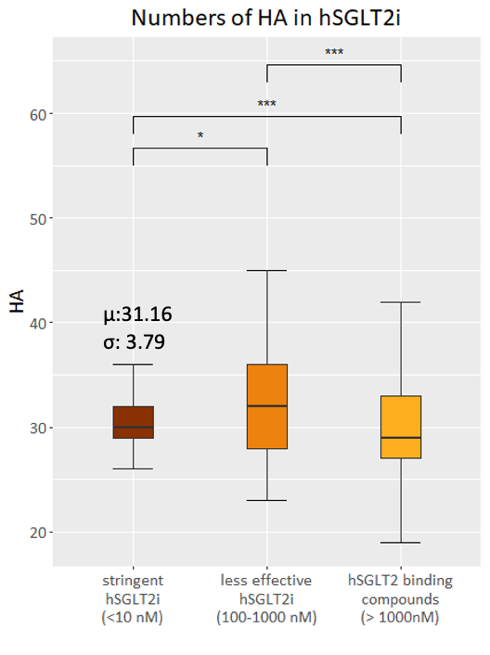

### Supplementary Data

**Supplementary 1.** HA distribution of hSGLT2i. The 692 hSGLT2i mentioned are composed by Less effective, intermediate, and stringent hSGLT2i (see Method). The HA interval for stringent hSGLT2i was determined within 24-38 (μ ±2σ).

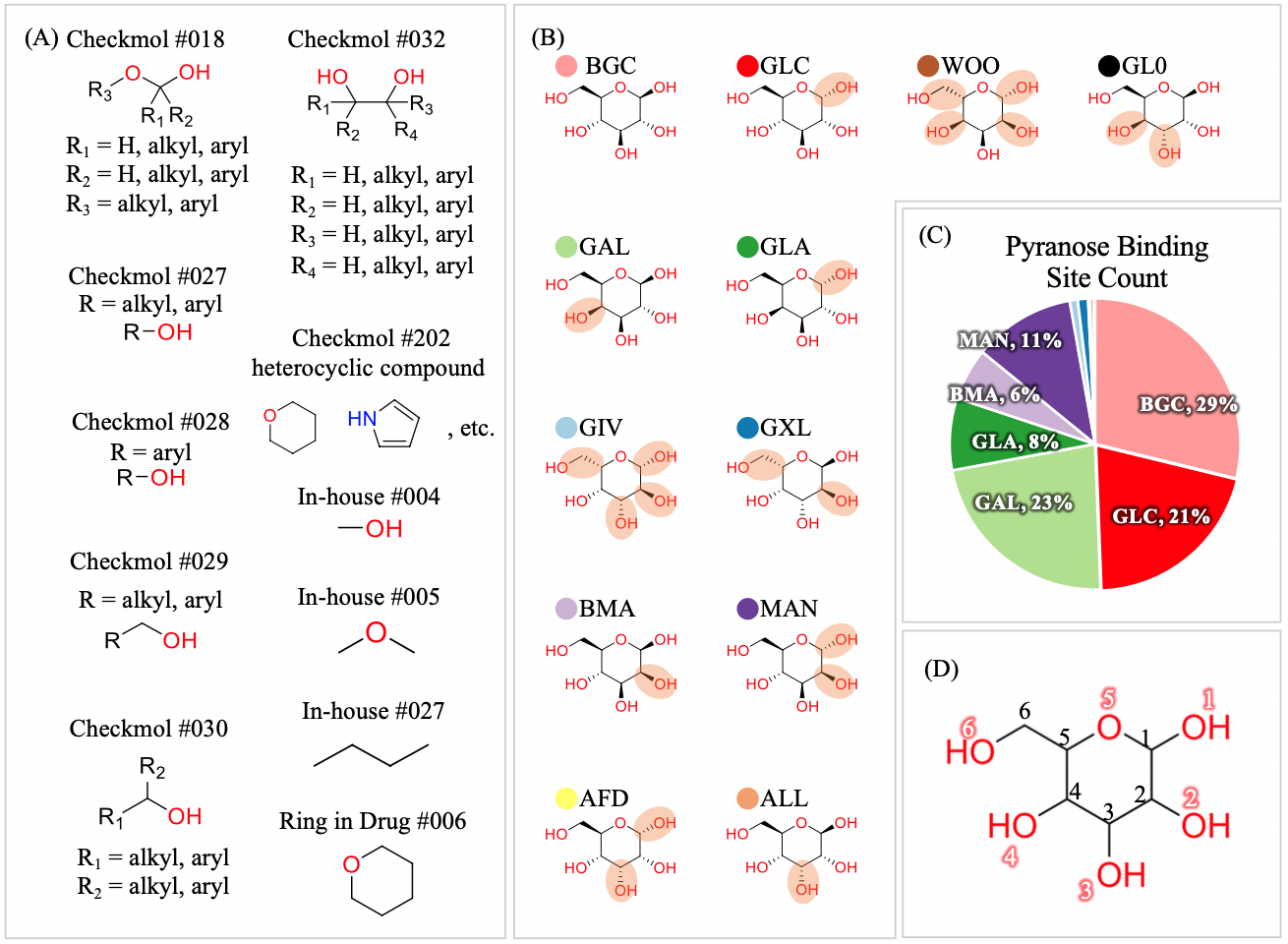
**Supplementary 2.** The composition of pyranoses and their protein binding sites. (A) The substructures dissected from pyranoses. (B) 12 of the 25 pyranoses with co-crystalized protein records in PDB. The orange circles highlight the stereo-difference comparing with BGC. (C) The distribution of pyranoses among 1572 binding sites. (D) The atom numbering of 25 pyranoses.

**Supplementary 3.** Propensity evaluates the specificity of a certain interaction pair in pyranose BSs. (A) Each interaction pair has a constant of preference in pyranose BS. Propensity of the pair is the logarithm of the given constant. 379 representative pyranose BSs are selected from set pyr (see Method) to establish the constant. Surface residues from Set b are taken as the general protein surface for further ratio calculation ((A)-1). The term is also normalized by the ratio of each considered atom in the pyranose compound ((A)-2). (B) An example of Q-O1 H-bonds in pyranose (PDB entry: 6h7d) and the process of establishing propensity under a given example of condition. The notation of 6h7d_A_BGC_600 refers to [PDB entry]_[protein chain]_[ligand ID]. (C) The propensity analysis of vdW on pyranose binding proteins. (D) The propensity analysis of H-bonds on pyranose binding proteins.

(-)

0

(+)

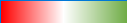

$$pro\left( Q,O1,H \right)$$

Frequents in pyranose interaction

Less seen in given
pyranose BSs

6h7d_A_BGC, … 379 BSs

x_pyr_(residue): GLN(Q)

j (atom type): O1

F (interaction): H-bond

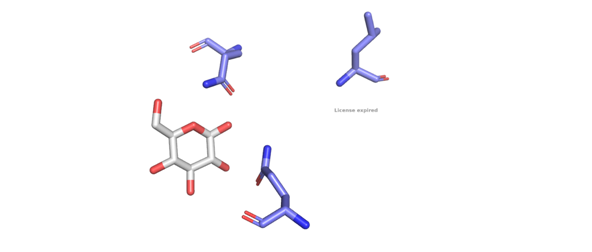

O1

Q177

Q295

(B)

-1.00

0

2.00

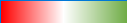

x (residue type in protein)

j (atom type in pyranose)

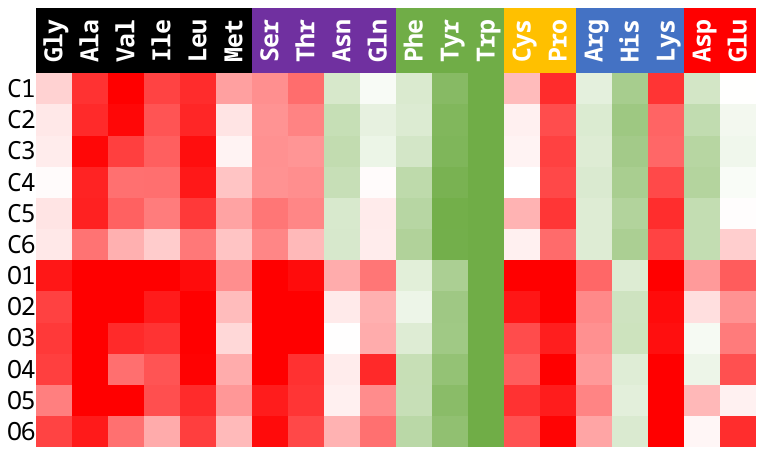

$$pro\left( x, j,vdW \right)$$

(C)

(D)

-1.00

0

1.50

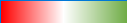

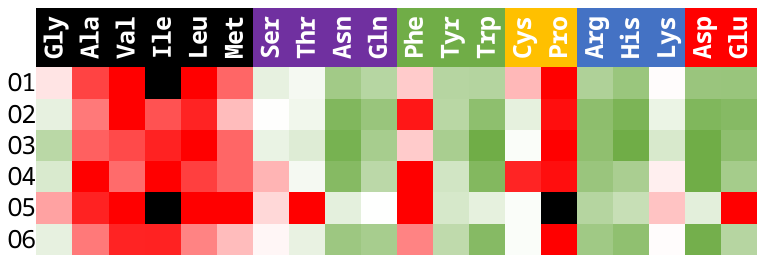

x

j

$$pro\left( x, j,H \right)$$

(A)-1

**Set b**

**Set pyr**

$$k_{xjF}=\frac{\frac{N_{x_{pyr}jF}}{\sum(N_{x_{pyr}jF})}}{\frac{n_{x_{b}}}{\sum n_{x_{b}}}\times\frac{n_{j}}{\sum n_{j}}}$$

(A)-2

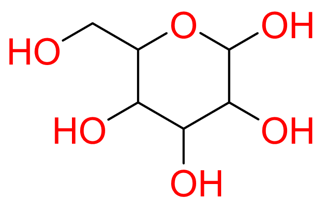

1

2

3

4

5

6

6

5

4

3

2

1

(A)

$$pro\left( x, j,F \right)=ln\left( k_{xjF} \right)$$

$$x:the given residue in one the 20 amino acids.$$

$$j:the atom type on pyranose, which can be O_{1}-O_{2}, or C_{1}-C_{6}$$

$$F:the contact to create the given force, H-bonds or vdW$$

**Supplementary 4.** Importance of O4 its differences with O1. (A) The H-bond 2D-diagram of hSGLT2-empagliflozin structure. Only the residues forming H-bonds to the compound is shown. The orange arrows point the atom corresponding to O1 and O4 in a pyranose. The plot was generated from LigPlot+ v.2.2.4. (B) The distribution analysis of H-bonds on each individual O in pyranose. The differences of the preference to form H-bonds among the Os show statistical significance (X^2^: 685.63, p < 2.2e-16). (C) The count of top 5 residues forming the most H-bonds to the O4 of pyranoses. Note that the top 3 residues are all double-headed polar on their side-chains.

H-bond distribution

on oxygen of pyranose

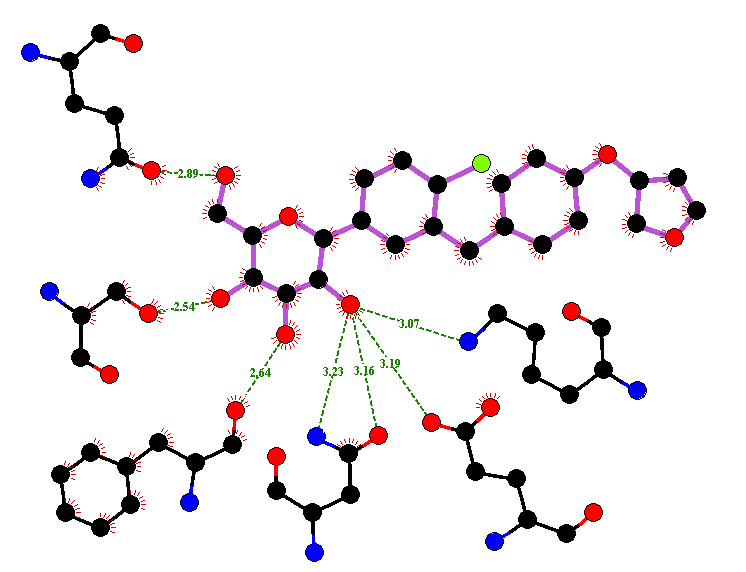

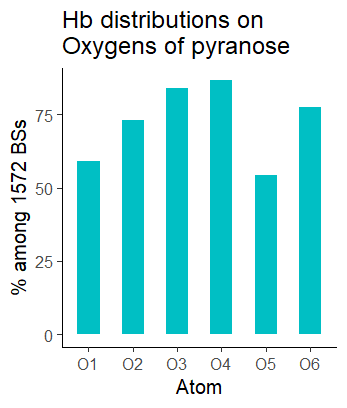

**hSGLT2 (7vsi)**

empagliflozin

C (O1)

O4

(A)

(B)

Counts of H-bonds to O4 of pyranoses

(C)

| Top 5residue |  |
| --- | --- |
| ASP | 749 |
| ASN | 326 |
| GLU | 305 |
| ARG | 194 |
| GLY | 147 |

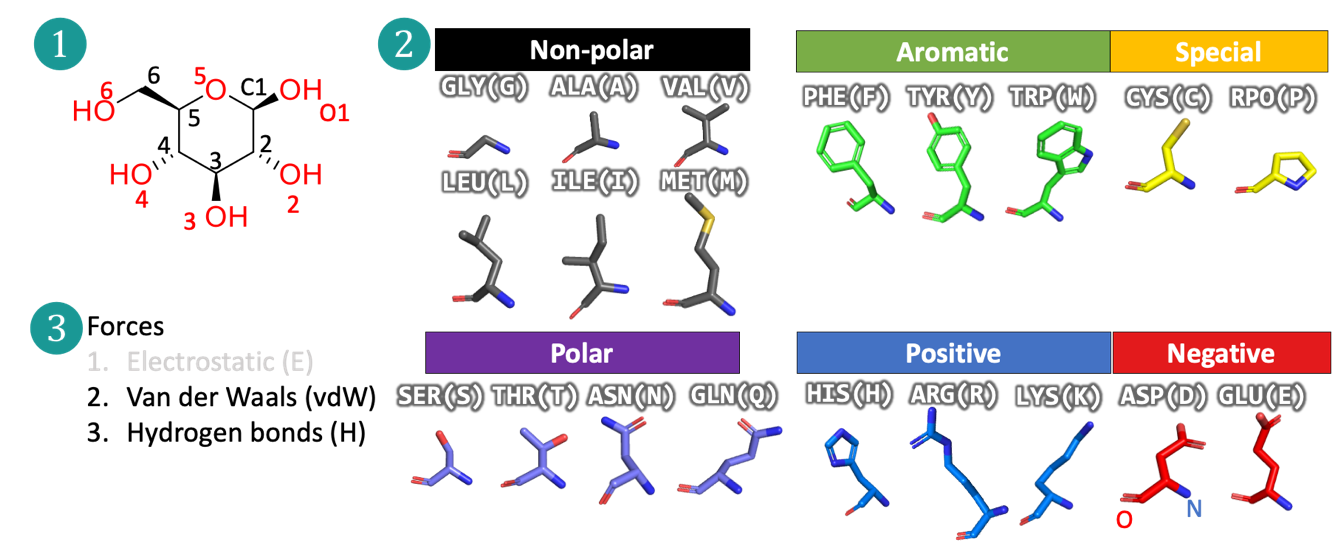
**Supplementary 5.** The composition of features in interaction profile of pyranose BSs.

| Glycine, GLY, G; | Alanine, ALA, A; | Valine, VAL, V; | Leucine, LEU, L; |
| --- | --- | --- | --- |
| Isoleucine, ILE, I; | Methionine, MET, M; | Serine, SER, S; | Threonine, THR, T; |
| Asparagine, ASN, N; | Glutamine, GLN, Q; | Phenylalanine, PHE, F; | Tyrosine, TYR, Y; |
| Tryptophan, TRP, W; | Cysteine, CYS, C; | Proline, PRO, P; | Histidine, HIS, H; |
| Arginine, ARG, R; | Lysine, LYS, K; | Aspartate, ASP, D; | Glutamate, GLU, E. |

**Supplementary 6.** The details determining pocket size. (A) The two major algorithms to evaluate the volume of a protein pocket (B) In cavity, SES display a larger pocket than SAS. (C) The yellow sticks connect the centers of alpha spheres composing a predicted pocket of fpocket. Red arrow points out the discontinuous spheres. The ligand is shown in white sticks. The protein is hidden in navy-blue background. (displayed with Swiss-PDBViewer^99^) (D)The two pictures are the same proportion of the same protein (PDB entry: 7dj2). fpocket does not adapt to the narrow mouth of the tunnel and sees the outer part as part of the predicted pocket. (visualized with PyMOL^48^) (E) When proteins are not parsed by chains (the right figure), the space in between are highly possible to be recognized as a binding pocket.

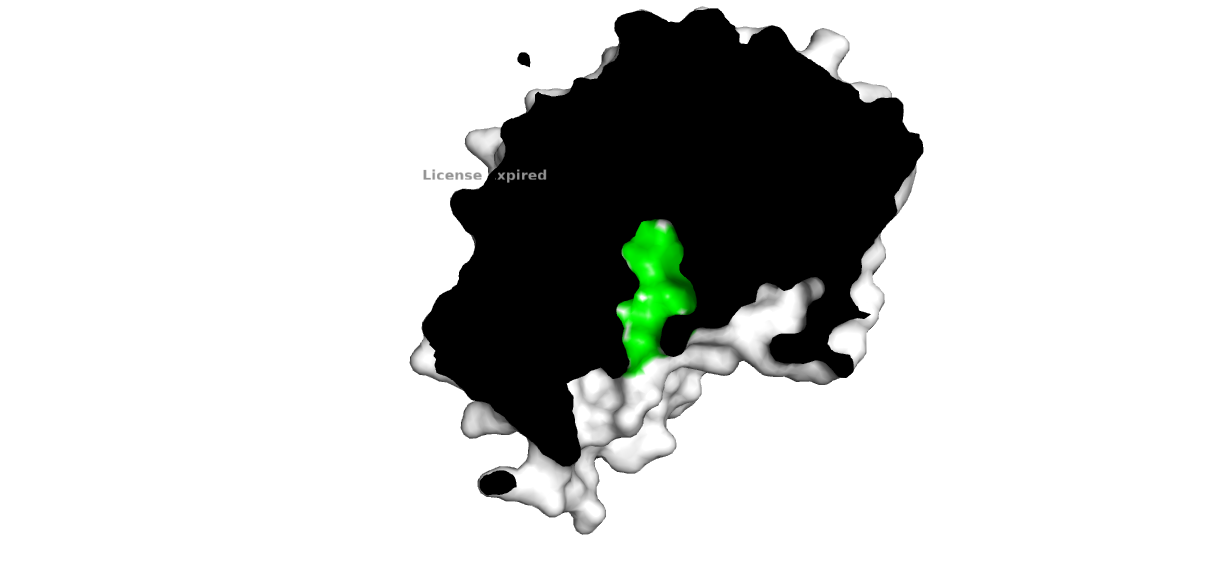

**6.4Å**

**CASTp**

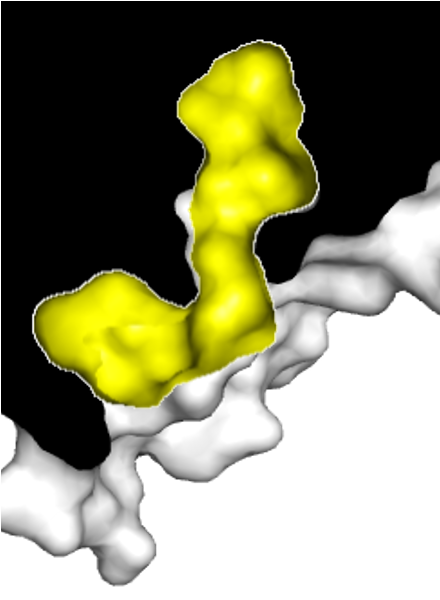

**fpocket**

**6.4Å**

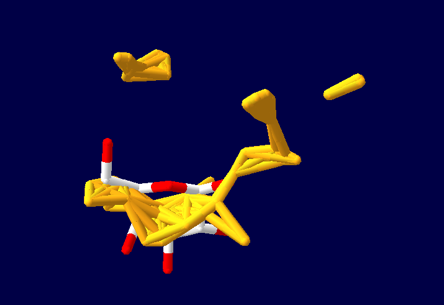

**fpocket**

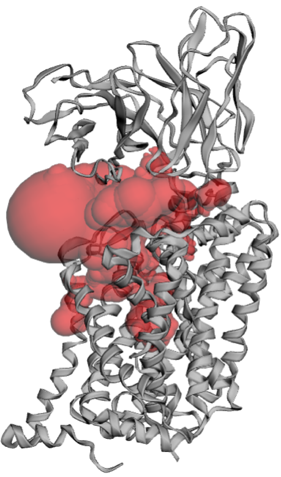

6vrk A chain
poc: 2335.595

6vkr whole protein

poc: 10059.15

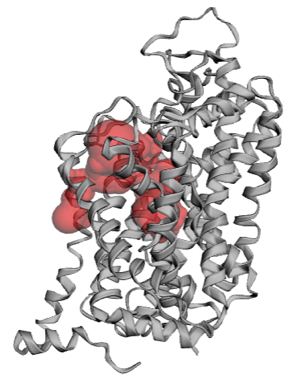

B chain

antibody

(E)

**CASTp**

(A)

(D)

(C)

Solvent accessible surface

vdW surface

Solvent excluded surface

(B)

Solvent molecules

atom

atom

atom

**Supplementary 7.** The surface residues count according to set b.

| **Amino acid** | **Whole** | **Main chain** | **Side chain** | **Amino acid** | **Whole** | **Main chain** | **Side chain** |
| --- | --- | --- | --- | --- | --- | --- | --- |
| GLY | 78,671 | 64,133 | 70,244 | PHE | 41,271 | 18,996 | 38,330 |
| ALA | 78,177 | 52,275 | 70,863 | TYR | 41,233 | 15,791 | 40,229 |
| VAL | 64,873 | 30,505 | 58,736 | TRP | 16,154 | 6,355 | 15,612 |
| ILE | 52,140 | 22,265 | 47,994 | CYS | 12,918 | 7,610 | 10,801 |
| LEU | 91,033 | 45,866 | 83,848 | PRO | 57,499 | 33,872 | 55,587 |
| MET | 18,979 | 9,778 | 17,879 | ARG | 70,519 | 34,494 | 70,283 |
| SER | 72,176 | 45,619 | 69,096 | HIS | 29,273 | 13,699 | 28,846 |
| THR | 63,918 | 31,518 | 61,536 | LYS | 79,682 | 44,887 | 79,546 |
| ASN | 53,858 | 30,903 | 53,037 | ASP | 74,879 | 43,380 | 74,140 |
| GLN | 49,973 | 25,997 | 49,572 | GLU | 92,823 | 52,654 | 92,462 |

**Supplementary 8.** The distribution of EA and pyranose pockets sizes. (A) EA serves as a primary filter for pocket size selection for pyranoses. The box plot was generated from 227/379 representative pyranose BSs forming a cavity at EA > 6. (B) Examples of pyranose BSs at different EA. The higher EA, the more compact the BSs. We finally determined that a protein pocket big enough to contain a pyranose should be > 180 Å^2^ as 2qvc.

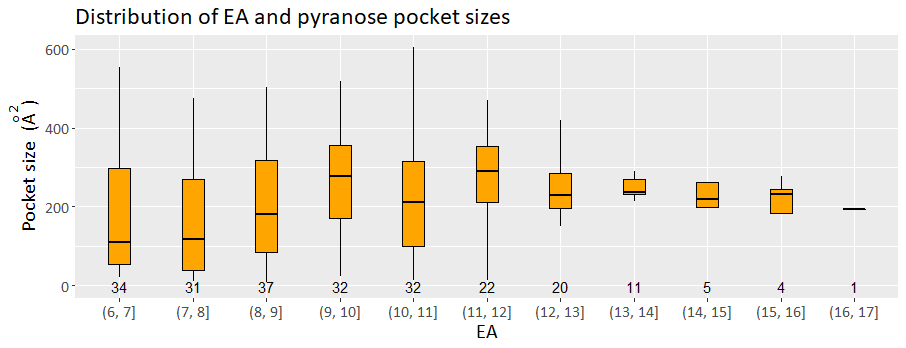

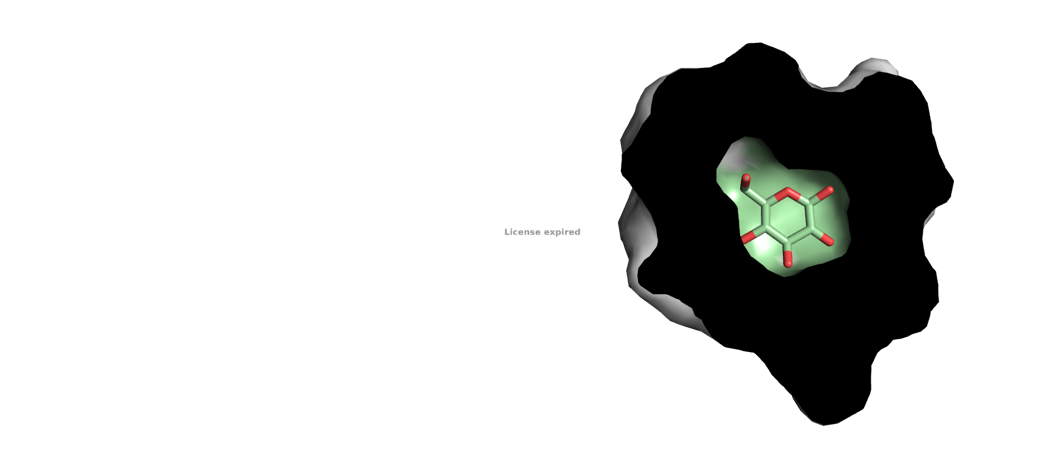

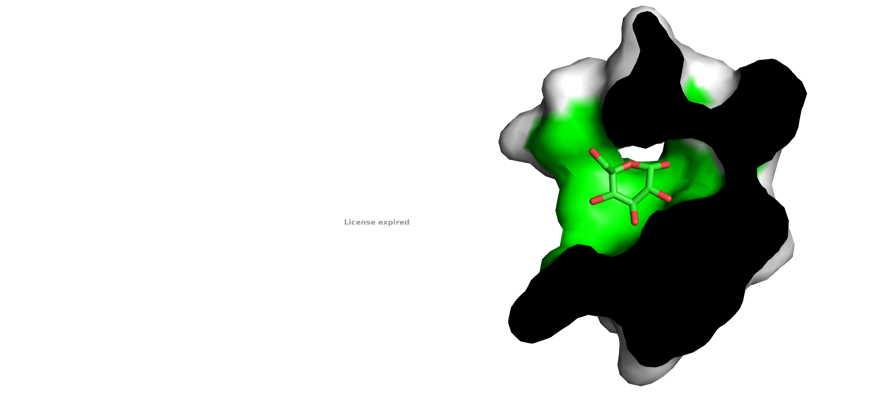

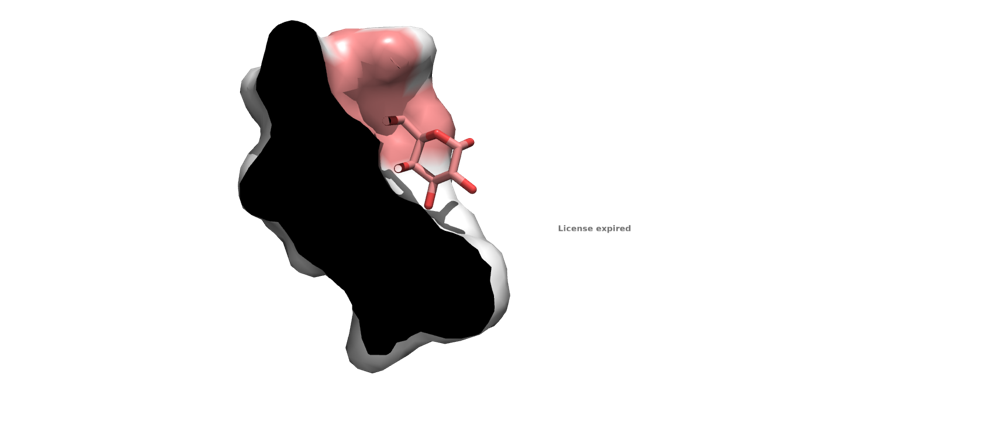

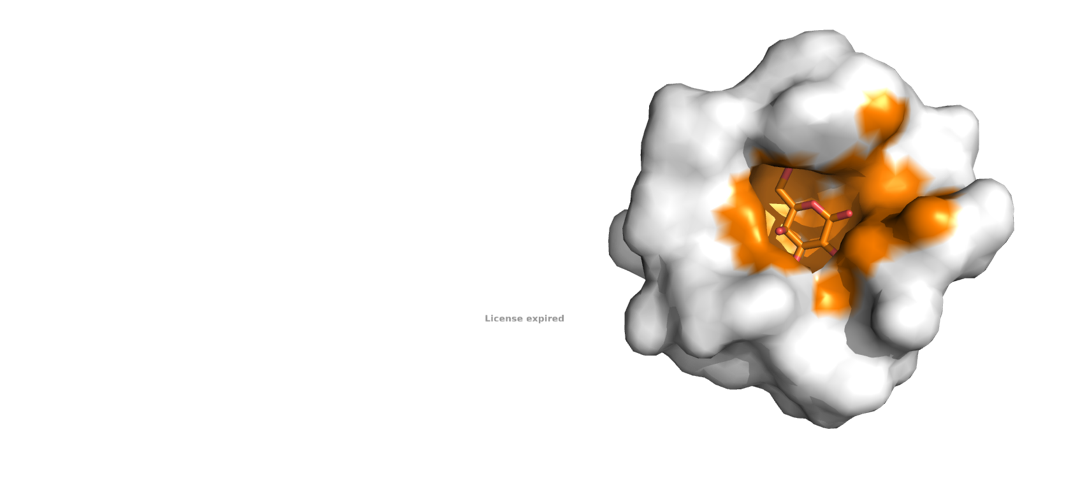

(B)

(A)

| **Protein** | **Agglutinin V**  **(plant)**  **(PDB entry: 3cal)** | **Bacterial xylose isomerase**  **(PDB entry: 4zbc)** | **Bacterial β-galactosidase**  **(PDB entry: 6qub)** | **Bacterial sugar transporter (PDB entry: 2qvc)** |
| --- | --- | --- | --- | --- |
| Pocket size (Å^2^) | 53.369 | 374.676 | 200.415 | 182.625 |
| 6Å-radius pocket |  |  |  |  |
